# Nervous system-wide analysis of all *C. elegans* cadherins reveals neuron-specific functions across multiple anatomical scales

**DOI:** 10.1101/2024.08.06.606874

**Authors:** Maryam Majeed, Chien-Po Liao, Oliver Hobert

## Abstract

Differential expression of cell adhesion proteins is a hallmark of cell type diversity across the animal kingdom. Gene family-wide characterization of their organismal expression and function are, however, lacking. We established an atlas of expression of the entire set of 12 cadherin gene family members in the nematode *C. elegans,* testing the hypothesis that members of this gene family define combinatorial expression codes that may drive nervous system assembly. This atlas reveals differential expression of genomically tagged cadherin genes across neuronal classes, a dichotomy between broadly- and narrowly-expressed cadherins, and several context-dependent temporal transitions in expression across development. Engineered mutant null alleles of cadherins were analyzed for defects in morphology, behavior, neuronal soma positions, neurite neighborhood topology and fasciculation, and localization of synapses in many parts of the nervous system. This analysis revealed a restricted pattern of neuronal differentiation defects at discrete subsets of anatomical scales, including a novel role of cadherins in experience-dependent electrical synapse formation. In total, our analysis results in novel perspectives on cadherin deployment and function.

## INTRODUCTION

The emergence of multicellularity was characterized by cell type diversification and accompanying elaboration of cell adhesion mechanisms (Hynes and Zhao 2000; Abedin and King 2008). among the most prominent families of cell adhesion molecules (cams) are the cadherins, large transmembrane proteins characterized by the presence of multiple protein-protein interaction domains, called cadherin repeats (takeichi 1988; Shapiro *et al*. 1995a; Shapiro *et al*. 1995b; Brasch *et al*. 2012; Brasch *et al*. 2018; Honig and Shapiro 2020). Based on primary sequence and/or the presence of additional domains, the cadherin superfamily encompasses multiple subfamilies - namely classical cadherin, protocadherin, calsyntenin, CELSR, dachsous, fat and fat-like, and cadherin-related (CDHR) - most of which are largely conserved across phyla (Nollet *et al*. 2000; Hulpiau and van Roy 2011; Nichols *et al*. 2012; Albertin *et al*. 2015). Cadherin family members orchestrate selective cell-cell recognition and communication events during developmental processes in many tissue types including the brain, ranging from cell migration, axon fasciculation and pathfinding, and synaptic connectivity (Hirano and Takeichi 2012; Loveless and Hardin 2012; De WIT AND GHOSH 2016; Yamagata *et al*. 2018; Sanes and Zipursky 2020). However, previous expression and functional studies on cadherins in animal brain development were restricted to a subset of cadherins, brain regions, neuronal classes, or developmental stages (Chen and Clandinin 2008; Schwabe *et al*. 2013; Duan *et al*. 2014; Schwabe *et al*. 2014; Mountoufaris *et al*. 2017; Duan *et al*. 2018; Tsai *et al*. 2020; Sun *et al*. 2021; Lv *et al*. 2022).

The genomic and cellular compactness of *C. elegans,* as well as its tremendous genetic toolkit make *C. elegans* an attractive choice to undertake expression and functional analyses of entire gene families, such as the cadherin family. Such comprehensive, gene family- and organism-wide analyses have the potential to reveal common themes in the deployment of specific gene families. For example, the expression and functional analysis of all members of the homeobox gene family revealed their re-iterative, combinatorial deployment in neuronal cell type specification throughout the entire nervous system of *C. elegans* (Reilly *et al*. 2020). Similarly, genome- and animal-wide analysis of neuropeptides and receptors has revealed intriguing novel perspectives on the abundance of neuropeptidergic communication in the nervous system (Ripoll-SANCHEZ *et al*. 2023) and the nervous system-wide expression analysis of classic neurotransmitter receptor families indicated the extensive and wide-spread use of non-synaptic neurotransmitter signaling in the nervous system (Yemini *et al*. 2021). Family-wide analyses of cell adhesion and recognition molecules offer the opportunity to tackle the fascinating problem of how neurons interact with one another to assemble into complex neuronal networks. Such network formation relies on the serial cellular interactions on different organizational and anatomical levels, ranging from cell migration and positioning to neurite outgrowth, selective fasciculation and synapse formation. One particularly intriguing question is whether evolution has selected individual gene families to be dedicated to any or all of these processes. What makes the cadherin family a particularly attractive candidate for a pervasively used determinant of brain patterning is the observation that, unlike for many other gene families involved in other critical neuronal processes (transcription factors, neuropeptides, ion channels), the number of cadherin genes scales with organismal complexity, suggesting a causal link between the two (Nichols *et al*. 2012). Moreover, the evolution of cadherin predates the origins of the nervous system (Abedin and King 2008), thereby prompting the hypothesis that cadherins may have had ancestral and, later, widely expanded functions in building neuronal networks.

The *C. elegans* genome encodes 12 cadherin genes (Hill *et al*. 2001), while mammals encode more than 100 members of the cadherin superfamily, a striking exemplification of how cadherin gene number scales with animal complexity (Nichols *et al*. 2012) (**Figure 1A**). While a small subset of the *C. elegans* cadherins have been previously analyzed (Broadbent and Pettitt 2002; Ikeda *et al*. 2008; Schmitz *et al*. 2008; Hoerndli *et al*. 2009; Steimel *et al*. 2010; Loveless and Hardin 2012; Najarro *et al*. 2012; Ohno *et al*. 2014; Sundararajan *et al*. 2014; Kim and Emmons 2017; Thapliyal *et al*. 2018a; Thapliyal *et al*. 2018b; Fan *et al*. 2019; Barnes *et al*. 2020; Hsu *et al*. 2020), the expression and function of several conserved cadherins (for example, the *C. elegans* homolog of Dachsous/DCHS) or the many non-conserved cadherins have not been previously examined at all. To explore how broadly cadherins are employed during the construction of the entire *C. elegans* nervous system, we first tagged all 12 *C. elegans* cadherins with a fluorescent reporter protein, using CRISPR/Cas9 genome engineering, and mapped their nervous system-wide expression in the developing and mature nervous system. We find that a subset of cadherins is expressed in all neuron types, while others are selectively expressed in a small subset of neurons. These patterns argue against a widespread use of combinatorial cadherin codes and rather suggest that cadherins fulfill either very broad generic themes in nervous system function (pan-neuronally expressed cadherins) or have very restricted, cell type-specific functions (narrowly expressed cadherins). To explore these possibilities further, we generated null alleles for all cadherins and extensively characterized them for defects in morphology, behavior, neuronal soma positioning, axonal neighborhood topology and fasciculation, axodendritic morphologies, and synapse localization to systematically decode overarching contributions of all cadherin members in establishing and maintaining neuronal structure. Our analysis reveals that different cadherin members play unique functions at various anatomical scales arising from several developmental events.

**Figure 1:**
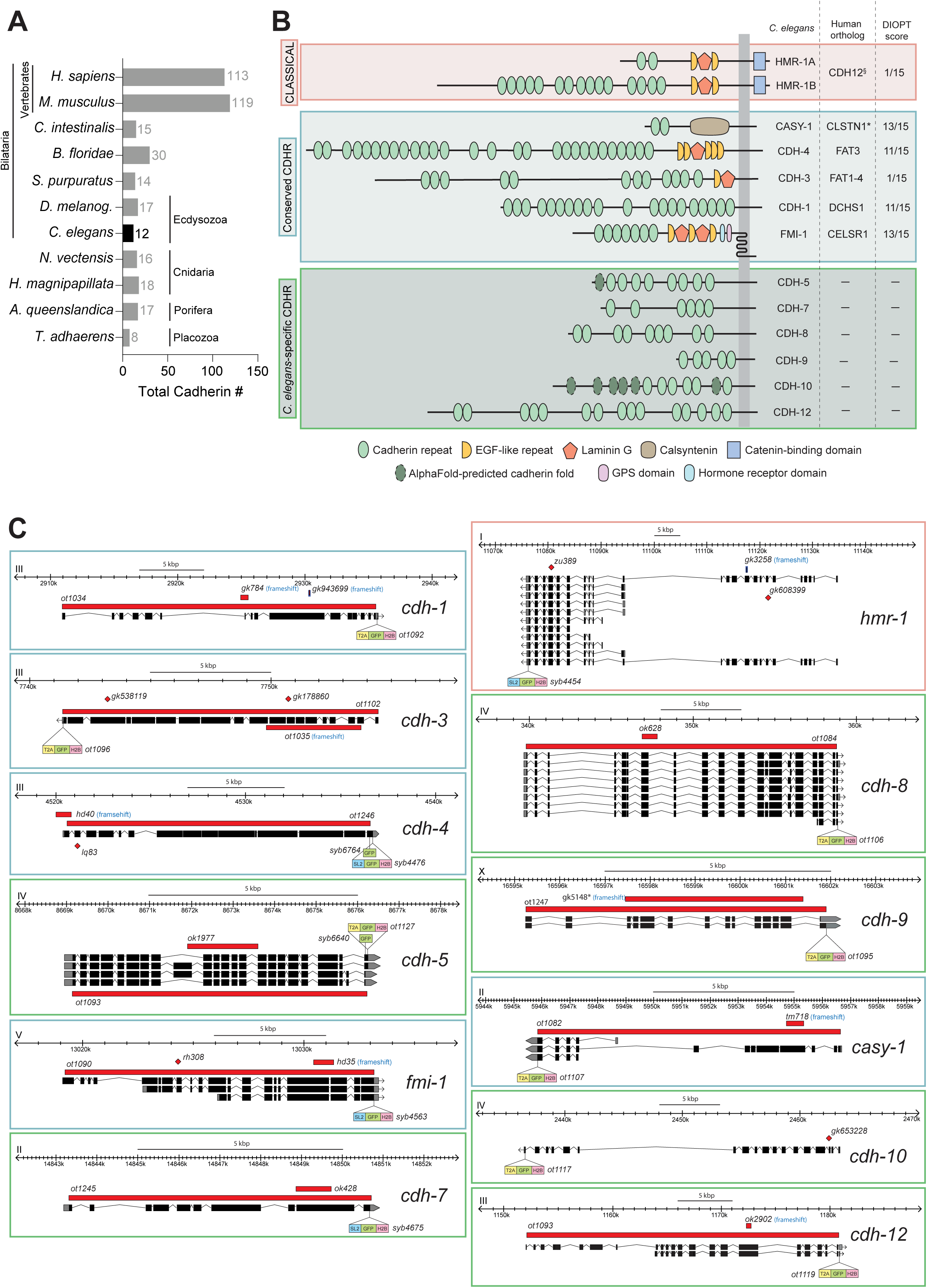
Overview of the cadherin superfamily in *C. elegans*. **(A)** Phylogenetic tree showing the number of cadherin-encoding genes expressed in marked species. *C. elegans* has 12 cadherin genes. **(B)** Schematic representation of *C. elegans* cadherin proteins. The *C. elegans* genome encodes 2 classical and 11 cadherin-related proteins, composed of cadherin repeats (light green ovals) and, in some cases, additional domains (legend). Closest vertebrate orthologs and DIOPT scores (Hu *et al*. 2011) are listed. Domain structure was retrieved from InterProScan (Quevillon *et al*. 2005); additional cadherin folds predicted using AlpaFold (Jumper *et al*. 2021) are shown (dashed dark green ovals). **(C)** Schematic representation of cadherin gene loci and CRISPR/Cas9 engineered reporter and mutant alleles. Each locus was endogenously tagged at the C-terminus with a T2A- or SL2-based GFP::H2B cassette to generate nuclear-localized reporters. In some cases, translational reporters were generated by tagging with GFP alone. Null alleles are depicted in red rectangles. Previous alleles, not definitively null, are also annotated.

## RESULTS

### A pan-Cadherin nervous system-wide expression atlas

The *C. elegans* genome codes for 12 members of the cadherin family, each characterized by the presence of multiple cadherin repeats (**Figure 1B**). Six of the cadherins are conserved across the animal kingdom and include the sole classic cadherin, HMR-1, the Fat orthologs CDH-3 and CDH-4, the Dachsous/DCHS ortholog CDH-1, the CELSR/Flamingo ortholog FMI-1 and the Calsyntenin/Clstn ortholog CASY-1, while another six appear to be nematode-specific, in line with frequent species-specific expansion of cadherin family members in other organisms (Nollet *et al*. 2000; Hulpiau and Van ROY 2011; Albertin *et al*. 2015). The protocadherin subfamily, characterized by the production of a vast array of isoforms (Mountoufaris *et al*. 2018), is absent in the genomes of *C. elegans* and there is generally only a very limited number of splice isoforms of *C. elegans* cadherins in general. While some of the conserved *C. elegans* cadherin genes have been characterized for expression and/or function in restricted parts of the nervous system (Broadbent and Pettitt 2002; Ikeda *et al*. 2008; Schmitz *et al*. 2008; Hoerndli *et al*. 2009; Steimel *et al*. 2010; Loveless and Hardin 2012; Najarro *et al*. 2012; Ohno *et al*. 2014; Sundararajan *et al*. 2014; Kim and Emmons 2017; Thapliyal *et al*. 2018a; Thapliyal *et al*. 2018b; Fan *et al*. 2019; Barnes *et al*. 2020; Hsu *et al*. 2020), none of these conserved cadherins have been comprehensively analyzed throughout the nervous system. In addition, none of the six nematode-specific cadherin proteins have been previously analyzed at all in terms of their nervous system expression or function.

We used CRISPR/Cas9 genome engineering to generate endogenously tagged GFP reporter alleles for all 12 *C. elegans* cadherin loci (**Figures 1C, 2A**). In each case, an SL2::GFP::H2B or T2A::GFP::H2B cassette was inserted at the C-terminus to capture expression of all putative splice isoforms, which all share common 3’ exons (**Figure 1C**). Because we expect many cadherins to display restricted subcellular expression patterns, including at synapses and neurites, we fused H2B to GFP to localize reporter signals to the nucleus, allowing us to unambiguously map the identity of cellular sites of expression using a polychromatic landmark reporter transgene, NeuroPAL (Neuronal Polychromatic Atlas of Landmarks) (Yemini *et al*. 2021) (**Figures 2A, S1A**).

**Figure 2:**
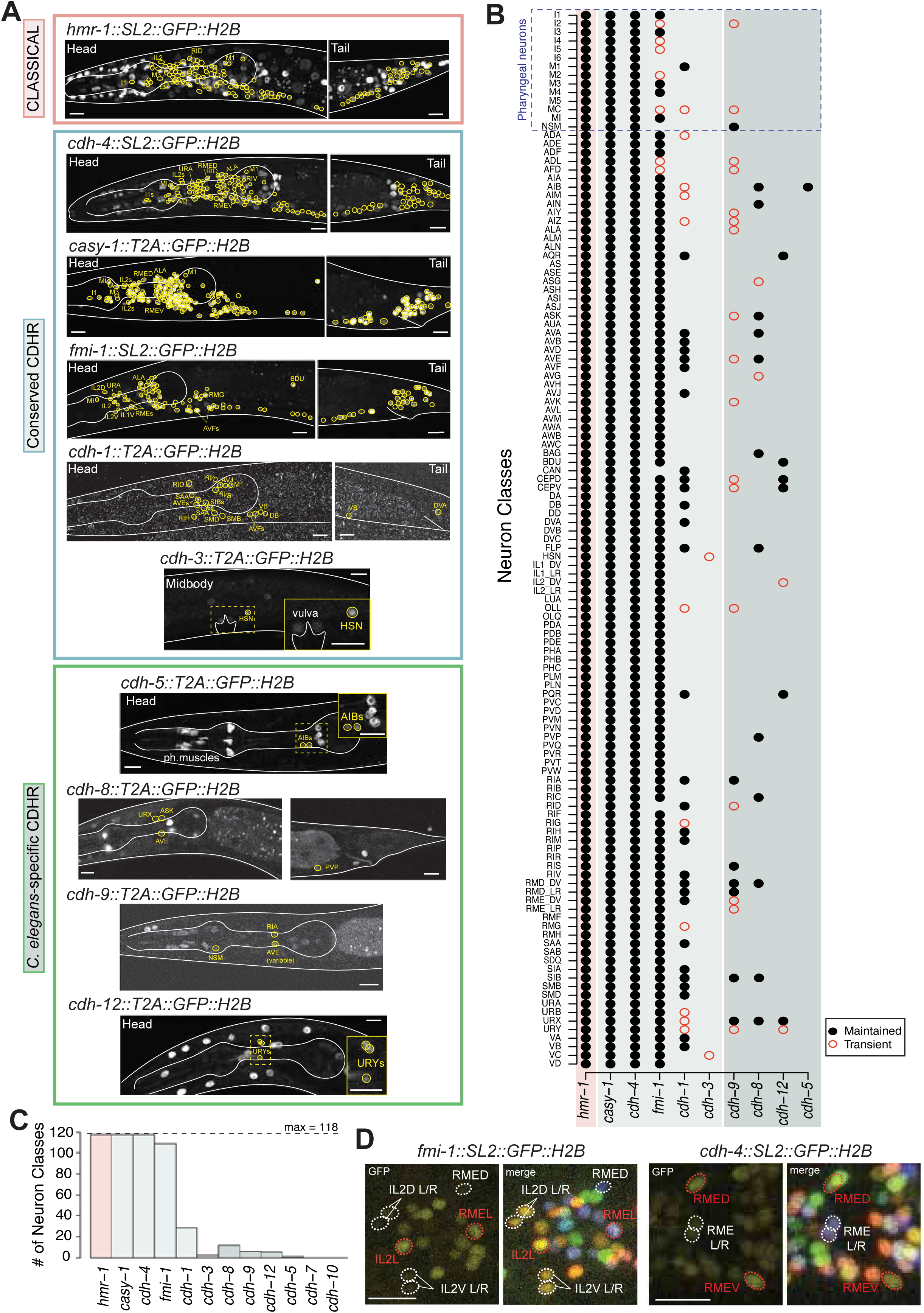
Cell-specific neuronal expression atlas of all cadherins. **(A)** Representative images of cadherin expression in the L4 nervous system. Reporters of classical cadherin *hmr-1*(*syb4454*) (pink) and other cadherins (green) *casy-1*(*ot1108*), *cdh-4*(*syb4476*), and *fmi-1*(*syb4563*) are pan-neuronally or broadly expressed. Reporters of cadherin-related members *cdh-1*(*ot1092*), *cdh-3*(*ot1096*), *cdh-5*(*ot1127*), *cdh-8*(*ot1106*), *cdh-9*(*ot1095*), and *cdh-12*(*ot1119*) are relatively narrowly expressed. All GFP+ neurons are encircled in yellow, but only some are labeled. **(B)** Nervous system-wide expression atlas in the L1 and L4 nervous system. GFP+ neurons were scored based on overlay with the NeuroPAL landmark reporter (Yemini *et al*. 2021) (*cdh-1* ID example in **Figure S1A**). Binary presence (black dot) or absence (no dot) of expression was scored. Transient expression is represented in a hollow red circle. **(C)** Abundance of each cadherin in the nervous system. **(D)** Subclass-specific differences in cadherin expression. *fmi-1*(*syb4563*) is enriched in IL2 L/R and RME L/R relative to IL2 D/V and RME D/V, respectively. In contrast, *cdh-4*(*syb4476*) is enriched in RME D/V relative to RME L/R. All images are of L4-stage animals and represent maximum intensity projections of a subset of the Z-stack. 5-10 animals were scored for each reporter. Worm body and the pharynx are outlined in white for visualization. Scale bar = 10μM.

We first mapped expression in the nervous system of late larval and young adult stages, when most of the nervous system has fully matured (**Figures 2B**). Overall, we found that cadherin expression patterns fall into two distinct categories: broad-versus narrowly-expressing (**Figure 2A-C**). Of the 12 cadherin genes, four were found to be expressed either pan-neuronally (*hmr-1*, *cdh-4/Fat*, *casy-1/Clstn*) or very broadly but not quite pan-neuronally (*fmi-1/Celsr*). Strikingly, all neuron types in which the *Celsr*/Flamingo homolog *fmi-1* is not expressed are located in the enteric (pharyngeal) nervous system of *C. elegans,* which has been proposed to constitute a more primitive domain of animal nervous systems (Furness and Stebbing 2018; Cook *et al*. 2020).

The remaining eight cadherins are narrowly expressed, but at varying levels of sparsity: four cadherins (the conserved *cdh-1/Dchs* and the nematode-specific *cdh-8*, *cdh-9*, and *cdh-12*) are expressed in 5-27 neuron classes (out of a total of 118 in the nervous system of the hermaphrodite), two are expressed in only 1-2 neuron classes (*cdh-3/Fat* in HSN and VC, and *cdh-5* in AIB), while the remaining two (*cdh-10* and *cdh-7*) do not show any detectable neuronal expression (**Figures 2A-C, S1B**). There is no correlation of neuron-specific cadherin expression with specific parts of the nervous system (sensory periphery, interneuron, motor circuit).

Expression levels of individual cadherins show stable differences across neuronal classes for each individual cadherin gene (**Figures 2A, S1C**). Expression differences can also be observed within members of the same neuronal class. For example, sensory neuron class IL2 and motor neuron class RME - both of which are further divided into subtypes based on anatomy and function - show differential cadherin expression across subtypes (**Figure 2D**) (Ward *et al*. 1975; White *et al*. 1986; Wang *et al*. 2010; Cros and Hobert 2022). The cadherin *fmi-1*/*Celsr* is enriched in IL2 L/R and RME L/R relative to IL2 D/V and RME D/V, respectively, and cadherin *cdh-4*/*Fat* is enriched in RME D/V relative to RME L/R. The *in vivo* relevance of quantitative expression differences in genes encoding CAMs - specifically cadherins - has been previously reported, such as in the case of Flamingo in *D. melanogaster* (Chen and Clandinin 2008).

Altogether, 61 of the 118 neuron classes of the late larval/adult nervous system do not express cell-type specific cadherins beyond the four broadly expressed cadherins *hmr-1, cdh-4, casy-1* and *fmi-1*. To frame this finding in the context of our original question of the expression specificity of the *C. elegans-*specific cadherins, we can conclude that cadherins do not form pervasive neuron-type specific codes of co-expression, at least in the mature nervous system.

There is no cadherin whose expression is restricted exclusively to the nervous system (summarized in **Figure 3L)**. Outside the nervous system, *hmr-1* is the most broadly expressed cadherin, even though not every cell expresses it (**Figure 3A**). Differential cadherin expression profiles in non-neuronal tissues include the hypodermis (the *C. elegans* epidermis), muscles, non-neuronal cells of the pharynx, seam cells, epithelial cells, and arcade cells (**Figure 3A-K**). Epidermal cells are clearly the most predominant site of non-neuronal cadherin expression, with *cdh-7* and *cdh-12* showing the greatest similarities in their broad epidermal expression (**Figure 3J, K**). Within the same type of non-neuronal tissue, cadherins are occasionally expressed in a subset of constituting cell types. The pharynx, the foregut of the worm which mediates food intake, is composed of a highly diverse set of cells, including 14 neuron classes and muscle, epithelial, gland, and marginal cells; in addition to neuronal expression (see above), we analyzed cadherin expression in all non-neuronal pharyngeal cell types and found notable patterns of cellular specificity in cadherin expression profiles (**Figure 3M**).

**Figure 3:**
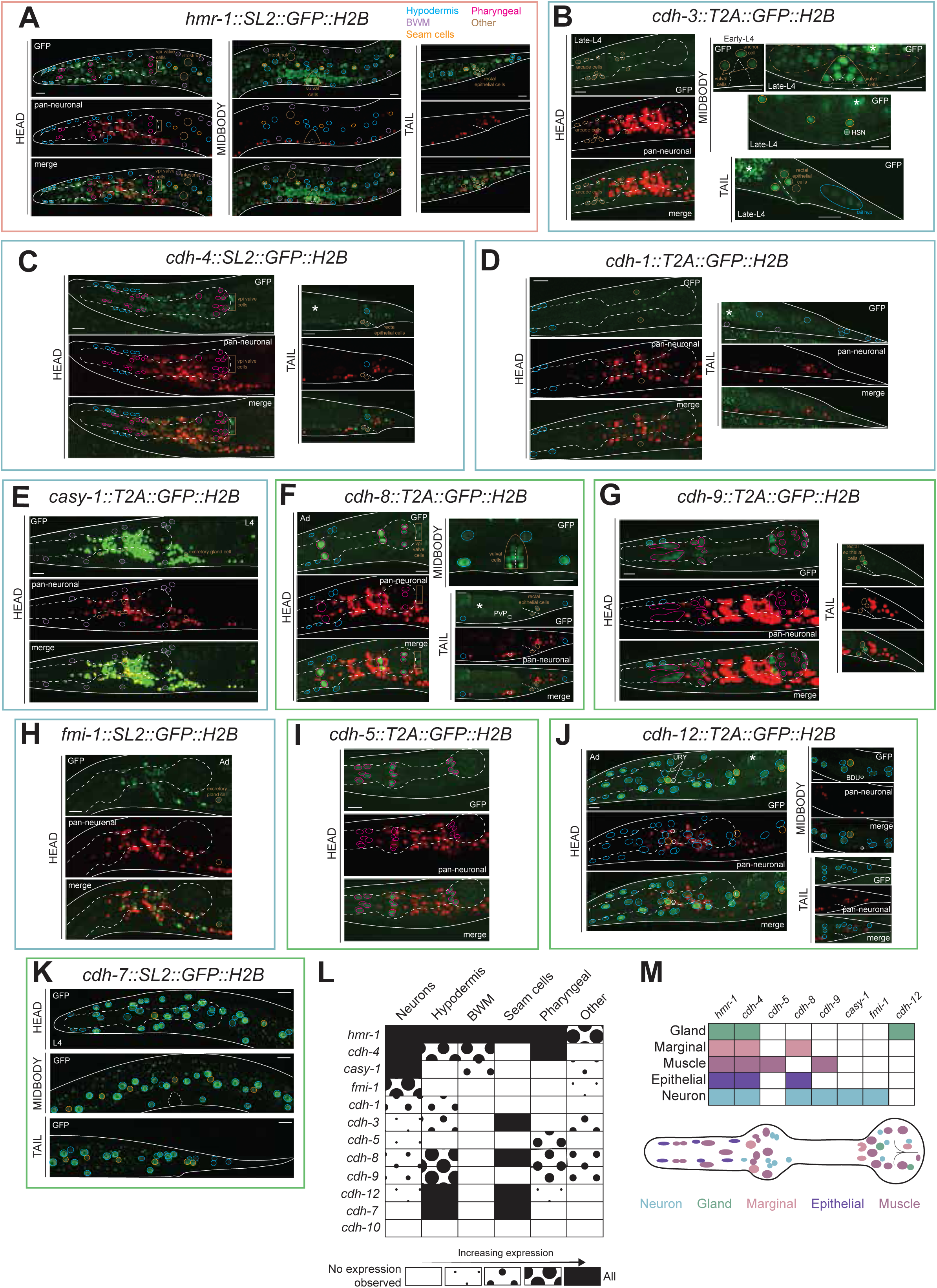
Cadherins are differentially expressed in non-neuronal tissues. Expression of cadherins in non-neuronal tissues, including the hypodermis (blue), muscles (purple), pharyngeal non-neuronal (pink), seam cells (orange), and others (brown). **(A)** *hmr-1* is expressed broadly expressed in all non-neuronal tissues. **(B)** *cdh-3* is expressed in vulval muscles, anchor cell, utse cell, seam cells, rectal epithelial cells, hypodermal cells in the tail, and arcade cells. **(C)** *cdh-4* is broadly expressed in multiple non-neuronal tissues, including hypodermal, pharyngeal, and rectal epithelial cells. **(D)** *cdh-1* is expressed in the hypodermis and seam cells. **(E)** *casy-1* is expressed in body wall muscles, although expression is significantly dimmer compared to neuronal expression. **(F)** *cdh-8* is expressed in pharyngeal, vulval, hypodermal cells, and the pharyngeal-intestinal valve (vpi). **(G)** *cdh-9* is expressed in various pharyngeal cells. **(H)** *fmi-1* is expressed in the excretory gland cell. **(I)** *cdh-5* is expressed in pharyngeal muscle cells. **(J-K)** *cdh-12* (J) and *cdh-7* (K) are expressed in hypodermal cells throughout the length of the worm. *cdh-7* is the only cadherin that is expressed exclusively in non-neuronal cells. **(L)** Summary of the extent of cadherin expression in various tissues, broadly categorized. Glial cells are excluded because glia-specific reporters have not yet been analyzed. **(M)** Cadherins are combinatorially expressed in non-neuronal cells of the pharynx. All images are of L4/Ad-stage animals and represent maximum intensity projections of a subset of the Z-stack. 5-10 animals were scored for each reporter. Gut autofluorescence is marked with an asterisk. Unidentified cells are circled in white. Scale bar = 10μM.

### Spatiotemporal changes in cadherin expression

Temporal changes in cadherin expression have been previously reported in many contexts (Kurusu *et al*. 2012; Mangione and Martin-BLANCO 2018; Knufer *et al*. 2020; Polanco *et al*. 2021). However, these studies were restricted to a small subset of cadherins, mostly classical cadherins, and were not performed with sufficient cellular specificity, on a nervous system-wide level, or across postnatal development. None of the *C. elegans* cadherins have been comprehensively assessed for expression dynamics with single neuron resolution. To fill these gaps, we sought to systematically assess changes in neuronal cadherin expression between developing and mature stages, with a focus on the developing nervous system.

We found that of the 12 cadherins, only two – the conserved classic cadherin *hmr-1* and the conserved *cdh-4/Fat* gene – are expressed early in embryogenesis, in dividing cells (**Figure 4A, B**); during this proliferative, pre-morphogenetic stages many short-range cell migratory events occur throughout the embryo, termed “cell focusing” (Bischoff and Schnabel 2006; Schnabel *et al*. 2006), as well as gastrulation. Early embryonic expression of these two cadherin genes correlates with *hmr-1/classical cadherin* and *cdh-4/Fat* mutants being the only cadherin mutants with prominent embryonic lethality, as we describe further below (Costa *et al*. 1998; Schmitz *et al*. 2008). The onset of all other cadherins is observed at about the time when most cells have started to exit the cell cycle and overall nervous system morphogenesis commences (**Figure 4A, B**).

**Figure 4:**
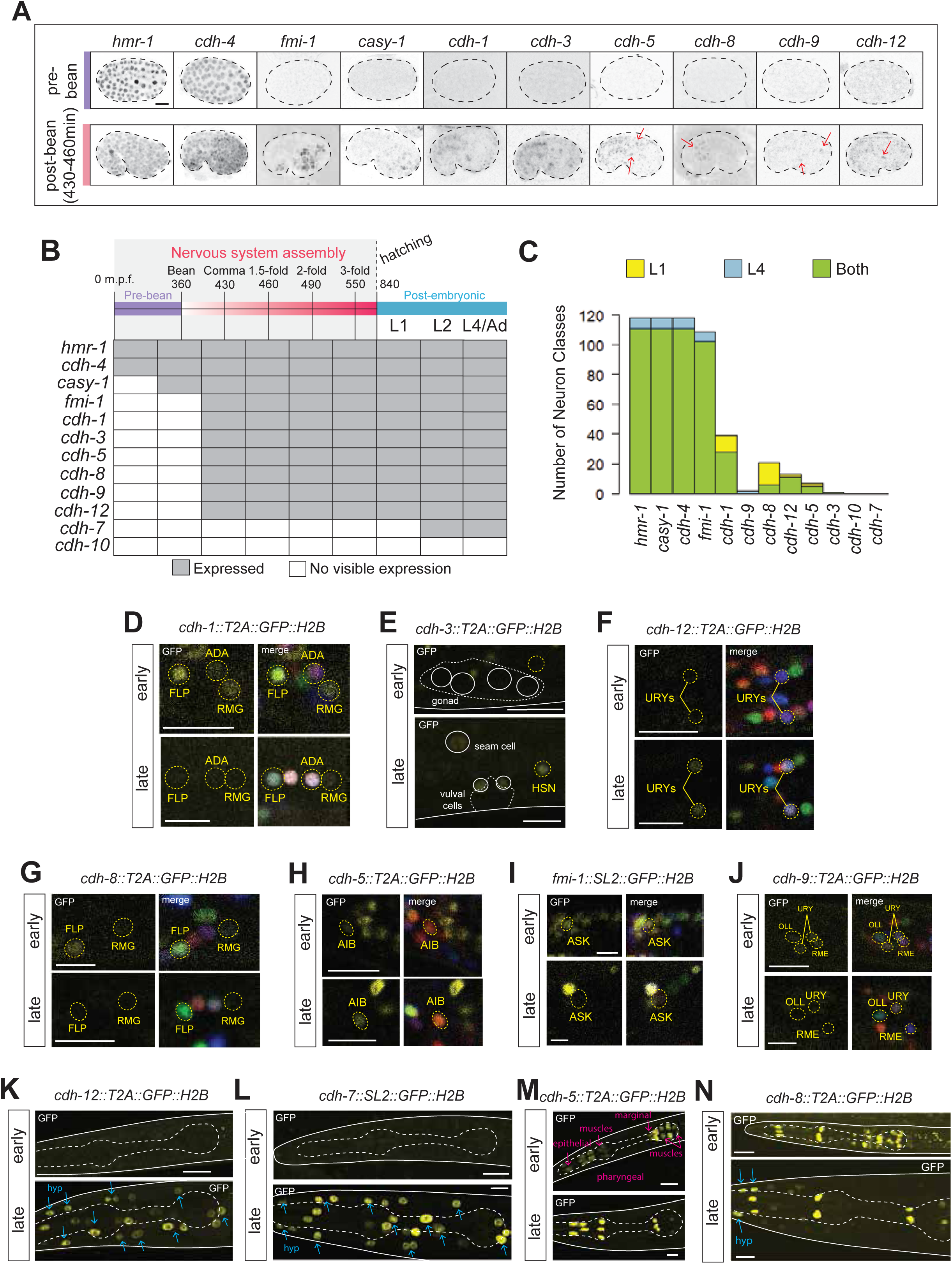
Widespread temporal changes in cadherin expression across development. **(A)** Differential onset of cadherin expression. All non-classical cadherins, besides *cdh-4*(*syb4476*), show detectable expression at the bean (*casy-1*) or post-bean (remaining cadherins shown) embryonic stage, and continue expressing at post-hatch (L1 stage); notably, onset of expression of cadherin-related genes coincides with neuronal differentiation and nervous system assembly. *cdh-7* is not expressed at any embryonic stage but starts expressing in non-neuronal tissues at the L2 larval stage (**Figure S1B**). No visible expression was observed for *cdh-10* at any stage in development (not shown), corroborating scRNA-seq data (Packer *et al*. 2019; Taylor *et al*. 2021). **(B)** Embryonic expression of all cadherin reporters in early and mid/late embryogenesis. **(C)** Comparative abundance of each cadherin in the nervous system of L1 and L4-stage animals. L1-only, L4-only, and shared expression is denoted in yellow, blue, and green, respectively. (**D-J**) Representative examples of temporal transitions in cadherin expression in the nervous system. Cadherins *cdh-1*(*ot1092*), *cdh-8*(*ot1106*), *fmi-1*(*syb4563*), and *cdh-9*(*ot1095*) are downregulated between early and late stages of development. Conversely, cadherins *cdh-3*(*ot1096*), *cdh-12*(*ot1119*), and *cdh-5*(*ot1127*) are upregulated. Expression changes are neuron-specific. (**K-N**) Representative examples of temporal transitions in cadherin expression in non-neuronal tissues. Cadherins *cdh-7*(*syb4675*) and *cdh-12*(*ot1119*) begin expressing in the hypodermis (blue) at the L2-stage. *cdh-5* shows restricted and enriched expression in the pharynx (pink) at late stage. *cdh-8* is upregulated in the hypodermis at late stage. All images are maximum intensity projections of a subset of the Z-stack. Gut autofluorescence is marked with an asterisk. Scale bar = 10μM.

Since the contorted shape and vigorous movement of late stage embryos in the eggshell severely limit the ability to precisely identify reporter gene expression patterns with reliable single cell resolution, we inferred cadherin expression at these critical stages by making use of the documented multi-hour (∼6h) stability of the H2B moiety in our reporter cassette (Dempsey *et al*. 2012). Such stability will lead a GFP::H2B reporter that turned on expression post-mitotically at mid/late embryogenesis to perdure until the initial hours of the first larval stage (L1). We therefore identified cadherin expression patterns at the L1 stage, with single neuron resolution using the NeuroPAL reporter, to infer cadherin expression at mid/late-embryonic stages when neuronal development and circuit formation occur (**Figures 2B, S2A**). The notion that expression patterns of H2B-based reporters at the first larval stage accurately reflect expression at earlier, mid-embryonic stages is further supported by two observations: 1) timing of cadherin expression onset visualized by our CRISPR reporters supports a previous embryonic single-cell RNA sequencing (scRNA-seq) dataset, and 2) our pan-cadherin expression atlas at the first larval stage and the embryonic scRNA-seq data show a similar dichotomy of broadly-versus narrowly-expressed cadherins, with largely congruent sites of expression (Packer *et al*. 2019).

Our L1 stage cadherin expression atlas leads us to conclude that cadherin expression is to some extent more widespread at developing stages compared to the L4/Ad nervous system. Specifically, we found that ten cadherins are expressed in the developing nervous system, five of which (*fmi-1/Celsr*, *cdh-1/Dchs*, *cdh-8*, *cdh-9*, and *cdh-12*) are more broadly expressed at the developing stage (**Figure 4C, S2A**). Of the remaining cadherins, three (*hmr-1*, *cdh-4/Fat*, and *casy-1/Clstn*) did not show significant changes in expression between developing and mature stages (**Figure 4C**). We did not find any detectable expression for cadherins *cdh-7* and *cdh-10* at the first larval stage (**Figure S1B**). While there is generally broader expression of cadherin expression during embryonic development of the nervous system, we consider it notable that 45 out of all embryonically generated neuron classes fail to express any cadherin other than the broadly expressed cadherins *hmr-1*, *cdh-4/Fat*, *casy-1/Clstn*, and *fmi-1/Celsr*. As stated above already in the context of the adult nervous system, we can conclude that cadherins do not form pervasive neuron-type specific codes of co-expression.

We found that changes in cadherin expression and directionality of change were gene- and neuron class-specific (**Figure 4D-J**). For instance, FLP sensory neurons express *cdh-1/Dchs* and *cdh-8* very highly at the developing stage but minimal or no expression, respectively, was detected at the late larval/early adult stages (**Figure 4D, G**). Similarly, *fmi-1/Celsr* is expressed more broadly at the developing stage compared to the mature stage (**Figure 4I**). While expression of most cadherins decreased between L1 and L4/adult stages, we observed an increase in expression in some cases. For example, *cdh-12* expression in URY sensory neurons increased across development (**Figure 4F**). *cdh-5* expression in AIB interneurons also increased across development but decreased in the late larval-to-adult transition, a window in which we found relatively few changes in cadherin expression (**Figures 4H, S2B**). Of note, when dynamic, expression did not necessarily change progressively in one direction. In the case of *cdh-5* in AIB neurons, for example, expression increased from L1 to L4, and then decreased in adults (**Figure S2B**).

Lastly, we observed additional changes in cadherin expression outside the nervous system (**Figure 4K-N**). The most striking change was in *cdh-7* and *cdh-12* expression; while no expression was observed at the L1 stage, both genes turned on their hypodermal expression at the start of larval stage L2, after many cells of the lateral hypodermis have stopped dividing (**Figure S2C**).

### Mechanisms of spatiotemporal regulation of cadherin expression

Many cadherins are expressed in a neuron type-specific manner, and thus contribute to transcriptomic diversity in the nervous system (**Figures 2, 4**). Extensive studies on the transcriptional regulation of neuron-specific terminal gene batteries have shown that terminal selector-type transcription factors are key regulators of postmitotic neuronal identities (Hobert 2016). While regulation of many gene families - including genes encoding CAMs such as immunoglobulin, neurotransmitter pathway, and other synaptic machinery molecules - has been pursued, cadherins are sparingly represented in these studies owing to a lack of good reporters and limited knowledge of expression patterns prior to this work (Berghoff *et al*. 2021; Cros and Hobert 2022; Vidal *et al*. 2022).

We asked if cadherin expression is controlled by terminal selector-type transcription factor mutants and indeed found this to be the case in several distinct cellular contexts. The terminal selector of AIB neuron identity, *unc-42* (Bhattacharya *et al*. 2019) controls *cdh-5* expression in AIB interneurons (**Figure 5A, F**)*. cdh-12* expression in the URY sensory neurons is lost in animals lacking the terminal selector of URY identity, the *unc-86* POU homeobox gene (SERRANO-SAIZ *et al*. 2013) and, similarly, expression of *fmi-1/Celsr* in IL2 sensory neurons, where *unc-86* also acts as a terminal selector (Shaham and Bargmann 2002; Pereira *et al*. 2019), is affected in *unc-86* mutants (**Figure 5B, C, F**).

**Figure 5:**
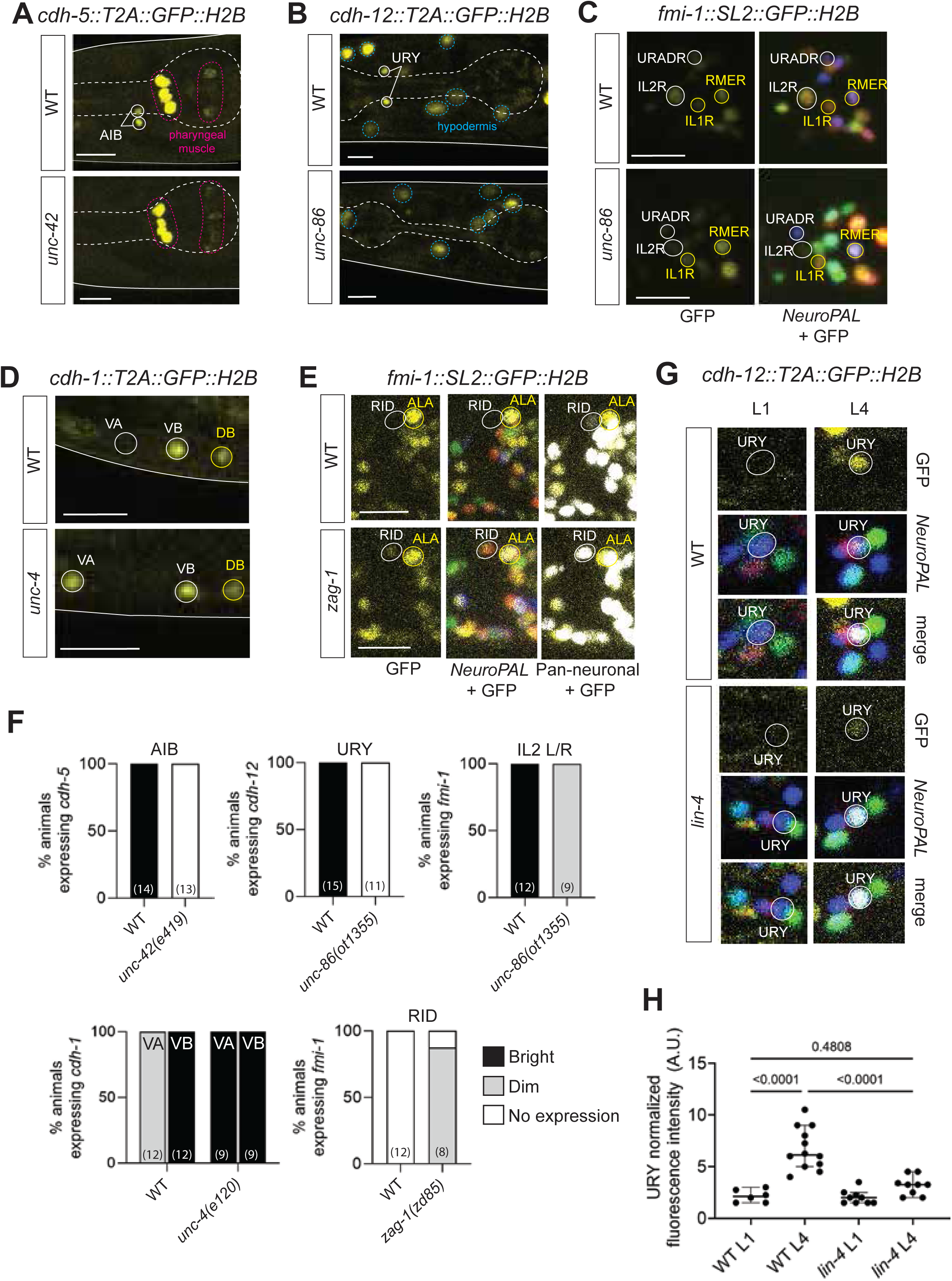
Spatiotemporal regulation of cadherin expression in the nervous system. (**A-E**) Representative images of cadherin reporters in wild type and mutant backgrounds. *cdh-5::T2A::GFP::H2B* (*ot1127*, AIB), *cdh-12::T2A::GFP::H2B* (*ot1119*, URY), and *fmi-1::SL2::GFP::H2B* (*syb4563*, only IL2R scored) are downregulated in mutants of terminal selector transcription factors UNC-42 (allele *e419*) and UNC-86 (*ot1355* for *cdh-12*, *ot1354* for *fmi-1*). *cdh-1::T2A::GFP::H2B* (*ot1092*) and *fmi-1(syb4563)* expression is derepressed in neurons VA and RID in transcription factors mutants *unc-4(e120)* and *zag-1(zd85)*, respectively. *cdh-1* and *fmi-1* expressing neurons were identified with NeuroPAL overlay. (**F**) Quantification of cadherin expression for (A-E). Statistical significance was assessed using the Fisher’s exact test or chi-squared test. (**G, H**) Representative images (G) and quantification (H) of *cdh-12*(*ot1119*) in wild type and *lin-4*(*e912*) mutant backgrounds in URY neurons. In wild type animals, *cdh-12* expression increases between L1 to late L4 stages. *cdh-12* expression in *lin-4* mutant L4 animals is significantly reduced and resembles L1 expression levels. All images are maximum intensity projections of a subset of the Z-stack. Scale bars = 10μM.

Motor neuron diversity in the ventral nerve cord of *C. elegans* is controlled by sets of repressor proteins that antagonize the function of the pan-cholinergic motor neuron selector *unc-3/EBF* (Kerk et al. 2017). We found that cadherins are controlled by a similar transcriptional regulatory logic. *cdh-1* - typically expressed in both lineally related sister neurons VA and VB but enriched in VB neurons - was derepressed in VA neurons in animals lacking the homeobox repressor *unc-4* (Miller et al. 1992), a known subtype selector that distinguishes VA from VB neurons (**Figure 5D, F**). Another broadly employed repressor of neuronal identity features, the *zag-1* Zn finger/homeobox gene (Clark and Chiu 2003; Wacker *et al*. 2003), also controls cadherin expression as we find that *fmi-1/Celsr* expression is derepressed in RID neurons in *zag-1* mutants (**Figure 5E, F**). We conclude that cadherins are substrates of a diverse set of transcriptional regulatory routines previously shown to generate spatial diversity of neuronal gene expression programs.

On the temporal axis, pathways that regulate changes in CAM gene expression are not well understood. Recent transcriptomic studies reveal the significant contribution of heterochronic pathway genes and ecdysone hormone signaling in mediating temporal expression changes in the postmitotic nervous system of *C. elegans* and *D. melanogaster*, respectively (Sun and Hobert 2021; Jain *et al*. 2022). We asked whether cell-specific temporal changes in cadherin expression were regulated by the heterochronic pathway. The heterochronic pathway comprises a cascade of miRNAs and proteins, including the miRNA *lin-4* that is required to downregulate the transcription factor LIN-14 to promote the temporal transition between early larval stages L1 and L2 (Ambros and Horvitz 1984). To assess whether temporal changes in cadherin expression are regulated by this pathway, we focused on *cdh-12*. In wild type animals, *cdh-12* expression in URY sensory neurons increases significantly between L1 and L4 developmental stages (**Figures 4F, 5G-H**). *lin-4* upregulation in the L1-L2 transition period being required to suppress juvenile and promote late-stage neuronal identity features and we indeed find that *cdh-12* expression is maintained in its juvenile “off stage” in *lin-4* mutants (**Figures 5G, H**). Thus, the expression of *cdh-12* in URY neurons is regulated by intersecting roles of the transcription factor UNC-86 and the heterochronic pathway on spatial and temporal axes, respectively.

### Mutant analysis of *C. elegans* cadherins

We used CRISPR/Cas9 genome engineering to generate mutant alleles of each individual cadherin locus (**Figure 1C**), except for the broadly and early expressed classical cadherin *hmr-1,* a previously well-characterized essential gene required for global embryonic morphogenesis (Broadbent and Pettitt 2002; Fan *et al*. 2019; Barnes *et al*. 2020) and the nematode-specific cadherins *cdh-7* and *cdh-10*, for which we detected no neuronal expression at any stage. For several of these 9 cadherins, no mutant allele has either existed or been examined before and for several others – including the conserved cadherins – previously used mutant alleles (mostly small deletions) are not unambiguous molecular null alleles. Hence, our strategy consisted of deleting the entire locus for each of the 9 cadherin genes.

We found that of all the 9 null mutant cadherin strains, only *cdh-4/Fat* null mutants display embryonic and larval lethality, albeit in an incompletely penetrant manner. As described with previous alleles, these embryonic and larval arrest phenotypes appear to be a consequence of severe morphogenetic defects (Steimel *et al*. 2010). Hence, the only two essential cadherins, *cdh-4/Fat* and *hmr-1,* that are required for embryonic development are those that are expressed throughout early and late embryonic development.

All other cadherin null alleles are viable, fertile and appear overall healthy. With the exception of *cdh-3/Fat* null mutants, which display tail morphogenetic defects as described previously (Pettitt *et al*. 1996) and *cdh-4/Fat* null animals that escape embryonic/larval arrest, no other cadherin mutants displayed obvious morphological defects. Again, except for *cdh-4/Fat* escapers, none of the cadherins display the types of obvious uncoordinated (Unc) locomotor defects that are characteristic for mutants of many genes involved in nervous system development or function. We subjected all mutants to a more detailed quantitative locomotory analysis using a MultiWorm tracker systems that measures multiple locomotory parameters. We observed defects in a subset of locomotory behaviors in *fmi-1/Celsr, cdh-4/Fat* and *cdh-5* null mutants but observed limited/no defects in animals lacking the pan-neuronally expressed, conserved *casy-1/Clstn, cdh-1/Dchs* or the nematode-specific *cdh-8 and cdh-9* cadherins (**Table S1**).

We undertook a more detailed mutant analysis of the two conserved, broadly neuronally expressed *cdh-4/Fat, fmi-1/Celsr* and more cell-type specifically expressed conserved *cdh-1/Dchs, cdh-3/Fat, as well* as the nematode-specific *cdh-5, cdh-8, cdh-9, and cdh-12* genes. Several aspects of *cdh-4/Fat* and *fmi-1/Celsr* had already been previously described and we broaden this analysis here further, while the other five cadherins had not been examined for nervous system development before. We excluded from this more detailed analysis the two panneuronally expressed cadherins whose function in nervous system development and function have been extensively described before, namely *hmr-1* (Broadbent and Pettitt 2002; Fan *et al*. 2019; Barnes *et al*. 2020) and *casy-1/Clstn* (Ikeda *et al*. 2008; Hoerndli *et al*. 2009; Ohno *et al*. 2014; Kim and Emmons 2017; Thapliyal *et al*. 2018a; Thapliyal *et al*. 2018b), and we excluded the non-neuronally expressed, nematode-specific *cdh-7*.

### The conserved cadherin *cdh-4/Fat* affects neuronal morphogenesis at multiple organizational levels

*cdh-4/Fat* null mutants display a partially penetrant embryonic or early larval arrest phenotype characterized by severe morphological malformations (Schmitz *et al*. 2008). Surviving animals display several neuronal phenotypes, including axon fasciculation and neuronal migration defects (Schmitz *et al*. 2008; Sundararajan *et al*. 2014). However, these previous analyses were restricted to a few neuron classes. To further expand mutant analyses across neuron types and anatomical structures, we next broadly assessed neuronal architecture – including soma positions, axon neighborhood placement, neurite contact, and synapse structure - in *cdh-4/Fat* mutants (**Figure 6**). *cdh-4/Fat* was previously implicated in neuronal migration in a subset of post-embryonically born midbody neurons that undergo long-range migrations (Schmitz *et al*. 2008; Sundararajan *et al*. 2014). We thus asked whether the migratory phenotype extended to most neuron classes that undergo short-range migration during embryogenesis. Neuronal soma positions were assessed using the fluorescent transgenic tool NeuroPAL (Yemini *et al*. 2021). Although positions were scored in the L4-stage nervous system for ease of scoring, our analysis considered the overall developmental history of cadherin expression in each neuron studied since we reasoned that a positioning defect induced at the embryonic or early post-embryonic stages would persist until maturity. We found that neurons located in the anterior ganglion had mispositioned soma in two independent *cdh-4/Fat* mutant alleles - our full deletion allele *ot1246* and allele *hd40* (Schmitz *et al*. 2008) (**Figures 6A, S3A**). Neurons were already mispositioned in L1 animals right after hatching, indicating that the phenotype is developmental rather than a failure to maintain initially correct neuronal positions until late larval/adult stages (**Figure 6A, 6B**). Neuronal defects in mice with mutations in the *cdh-4* homolog *Fat* arise from over-proliferation of progenitors and their failure to differentiate into neurons (Cappello *et al*. 2013), but we did not find similar phenotypes in our NeuroPAL analysis: *cdh-4(ot1246)* mutants and wild type animals had the same number of neurons in the head, assessed based on a synthetic pan-neuronal reporter included in the NeuroPAL strain (**Figure S3B**).

**Figure 6:**
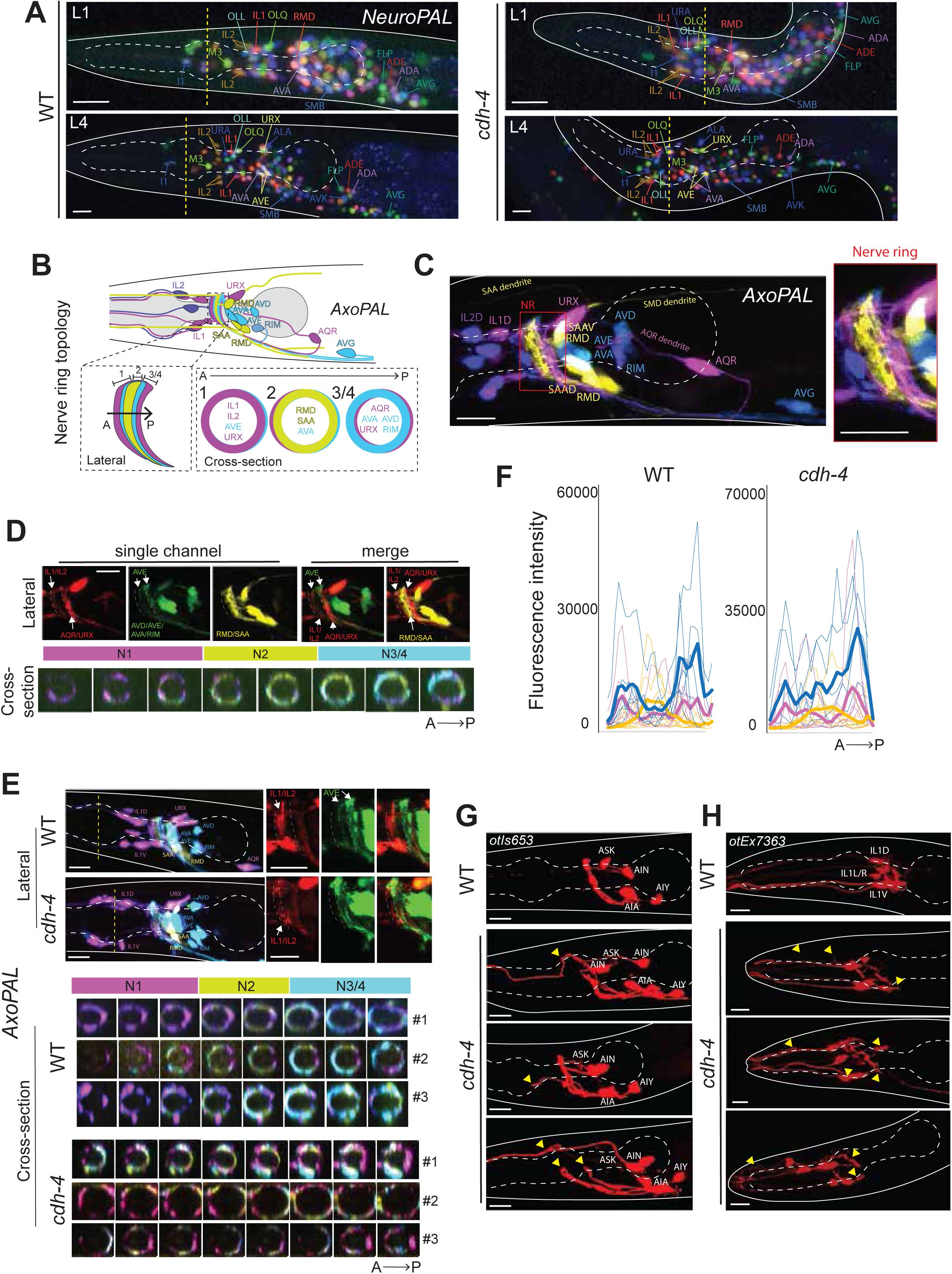
Broad phenotypic characterization of *cdh-4/Fat* mutants. (**A**) Neuronal soma mispositioning defects in *cdh-4(ot1246)* mutants. Representative images of soma positions in the landmark reporter NeuroPAL (*otIs669*) are shown. For reference, the middle of the anterior pharyngeal bulb is marked with a yellow dashed line. **(B, C)** AxoPAL reporter design for assessing axonal topology in the nerve ring. Schematic illustration (B) and a representative image (C) of AxoPAL reporter (*otEx7895*). A subset of neuron classes is cytoplasmically labeled with four distinct fluorophores to visualize 3 distinct neighborhoods in the nerve ring: N1 (magenta), N2 (yellow), and N3/4 (cyan) along the anteroposterior axis. **(D)** Lateral and cross-sectional representations of AxoPAL to depict positioning of labeled neurons in distinct axonal neighborhoods. IL1 and IL2 occupy the anterior-most neighborhood, N1. Neurons AVD, AVE, AVA, RIM, AQR, and URX largely occupy N3/4; the neighborhood shift of the command interneuron AVE is visible. Neurons RMD and SAA occupy neighborhood N2, which is mutually exclusive from N1 and N3/4. **(E)** *cdh-4(ot1246)* mutants have variable defects in axonal topology. Representative images of lateral and cross-sectional views of AxoPAL are shown. **(F)** Neighborhood-constituent reporter intensity traces along the A-P axis in *cdh-4(ot1246)* and WT animals. Bold line graphs represent an average of multiple worms. **(G, H)** Variable axodendritic defects in AIN (H) and IL1 (I) neurons. All images are maximum intensity projections of a subset of the Z-stack. 5-10 animals per genotype were analyzed for the NeuroPAL and AxoPAL analyses. Scale bars = 10μM.

Given prior evidence of cell body positions being correlated with layered nerve ring axonal topology (Moyle *et al*. 2021), we reasoned that defects in neuronal soma positioning in *cdh-4/Fat* mutants would in turn perturb nerve ring organization. Typically, such analysis would rely on multiple reporters that fluorescently label single or very sparse subsets of neurons; this is tedious and precludes analyzing relative axon positions in process-dense neuropils such as the *C. elegans* nerve ring. Conversely, multicolor reporters that selectively label neuronal subsets not only expedite analysis but provide relative spatial information (Hutter 2003). We therefore engineered a multicolor axonal reporter, which we term “AxoPAL”, that cytoplasmically labels a subset of head neurons (IL2, IL1, URX, AVA, AVE, AVD, RIM, AVG, RMD, SAA, and AQR) in four distinct colors to be able to spatially disambiguate their nerve ring axonal placement (**Figure 6B, C**). Briefly, AxoPAL resolves three distinct nerve ring neighborhoods (N1, N2, and N3/4 from left to right along the AP axis) both laterally and cross-sectionally (**Figure 6D**). Anterior ganglion inner labial neurons (IL1, IL2) project axons posteriorly to mainly occupy N1 and loop across all posterior neighborhoods. Neurons in ganglia posterior to the nerve ring project anteriorly; these include RMD and SAA (N2, yellow), and AVA, AVE, AVD, RIM, AQR, and URX (N3/4). AVE, URX, IL1, and IL2 discernibly occupy multiple neighborhoods in AxoPAL, which is consistent with previous EM-based studies (**Figure 6D**) (White *et al*. 1986; Brittin *et al*. 2021; Moyle *et al*. 2021). Unlike the nuclear reporter NeuroPAL, AxoPAL labels only a subset of neurons to achieve sufficient spatial resolution to visualize neuronal processes. Importantly, AxoPAL-labeled neurons were chosen based on each class expressing a subset of narrowly expressing cadherins at one or multiple timepoints in development (**Figure 2B**).

In *cdh-4/Fat* mutants, we found that the neuron-specific axonal composition in each neighborhood was disrupted, such that distinctive neighborhoods failed to be established, with the most obvious disorganization defects in interneurons (AVA/AVE/AVD/RIM labeled in cyan); in addition, the AQR axon was missing in most animals since the cell body fails to migrate anteriorly in the absence of *cdh-4/Fat* as shown previously (Schmitz *et al*. 2008) (**Figure 6E, F**). Apart from disrupted neighborhood topology, the nerve ring was anteriorly shifted in a subset of *cdh-4/Fat* mutant animals (**Figure S3C**). On the macroscopic organizational level, we also observed variable defects in axodendritic morphologies – as shown for neuron classes AIN and IL1 – in *cdh-4* mutants (**Figure 6G, H**).

### Effects of *cdh-4/Fat* on synaptic structure

Extending our analysis of neuronal anatomy to finer structures, we next asked whether loss of CDH-4/Fat affects synaptic structure. Here we took advantage of a recently developed repertoire of synaptic reporters comprising two types of reporters: 1) reporters that visualize all presynaptic specializations of a neuron via fluorescently-tagged synaptic proteins such as CLA-1/Piccolo or RAB-3 to label all synapses made by that neuron, and 2) reporters that use split-fluorophore technology, such as GRASP and iBLINC, to specifically label synapses between two neuron types (Feinberg *et al*. 2008; Desbois *et al*. 2015; Majeed *et al*. 2024).

First, we studied the role of *cdh-4/Fat* in the context of PHB and AVA synapses. We analyzed multiple features of PHB and AVA development to assess if they are dissociable. *cdh-4/Fat* null mutants did not show a significant PHB/AVA soma positioning defect (**Figure 7A-D**) and the number of synapses between PHB and AVA – assessed with an NLG-1 GRASP reporter – was also unaffected at both L1 and L4 stages (**Figure 7E, F**). We observed that PHB>AVA synapses in *cdh-4/Fat* mutants were clustered in regions of PHB/AVA fasciculation and excluded where PHB and AVA axons lost contact (**Figure 7E**). This prompted us to ask if CDH-4/Fat’s role in mediating PHB/AVA neurite contact and synapse formation could be dissociated. We found that PHB-AVA neurite contact - visualized with a CD4-GRASP reporter that we previously engineered (Liao *et al*. 2024) – was significantly reduced in *cdh-4/Fat* mutants; again, this phenotype was present at both L1 and L4 stages, suggesting that it is not a defect in the maintenance of neurite contact, rather the establishment of contact is affected early in development (**Figure 7G, H**). Our results suggest that CDH-4/Fat mediates contact between PHB and AVA but is dispensable for synapse formation.

**Figure 7:**
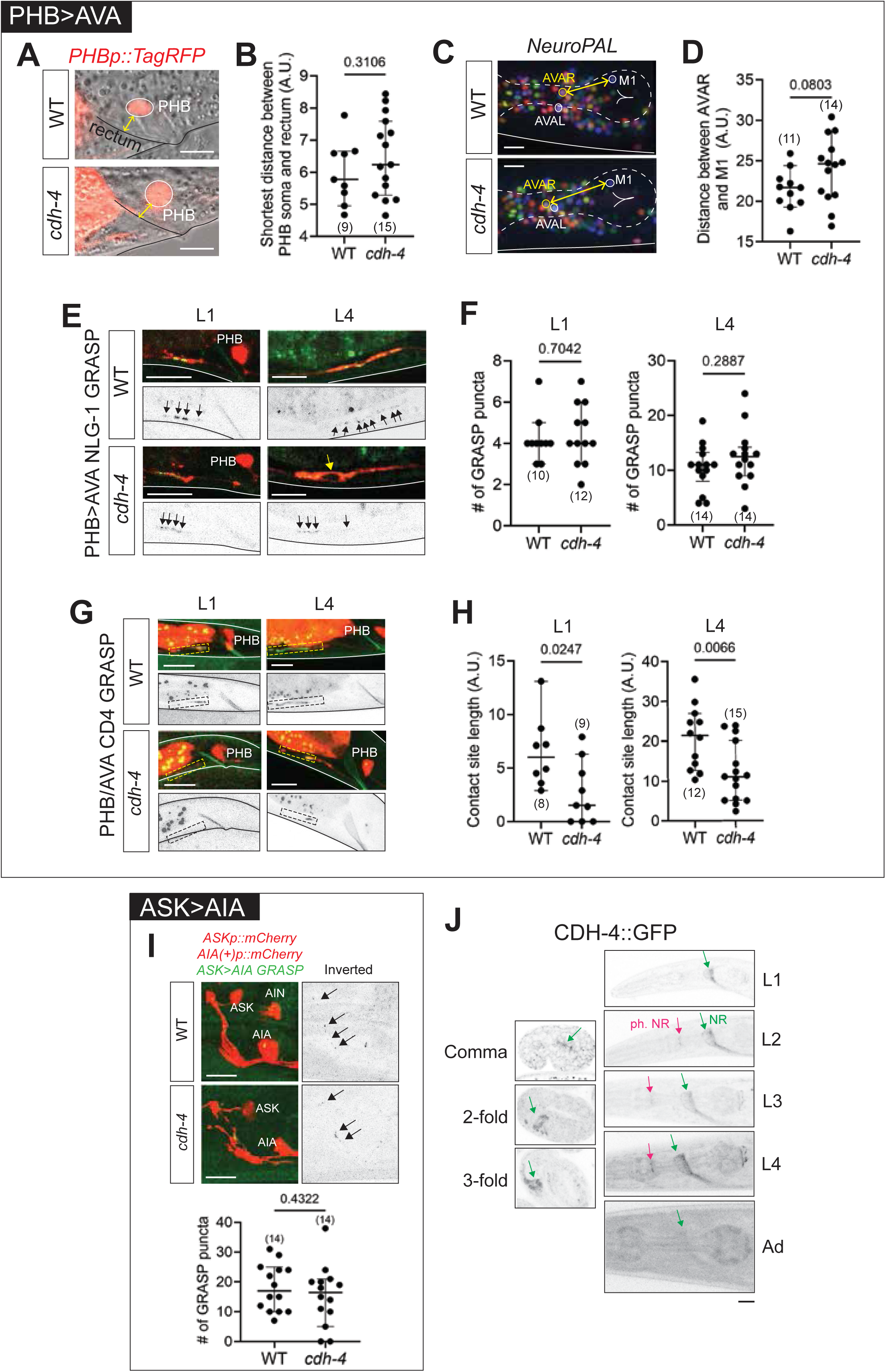
*cdh-4* mutants have context-specific effects on neuronal soma positions, axonal fasciculation, and synapse formation. (**A, B**) PHB soma positions are unaffected in *cdh-4* null mutants. Representative images (A) and quantification (B) of PHB soma labeled with *srab-20p::TagRFP* (*otEx8152*) in wild type and *cdh-4(ot1246)* null mutant animals. The shortest distance (yellow) between PHB soma and the rectum was quantified to assess PHB soma position in L4 stage animals. (**C, D**) AVA soma positions are not affected *cdh-4* null mutants. Representative images (C) and quantification (D) of AVAR soma labeled with *NeuroPAL* (*otIs669*) in wild type and *cdh-4(ot1246)* null mutant animals at the L4 stage. The shortest distance (yellow) between AVAR and M1 soma, both in the same z-plane, was quantified to assess AVA soma position. M1 is unaffected in *cdh-4* null mutants. (**E, F**) PHB>AVA synapses are unaffected in *cdh-4* null mutants. Representative images (E) and quantification (F) of PHB>AVA NLG-1 GRASP (*otIs839*) puncta in wild type and *cdh-4(ot1353)* null mutant animals at L1 and L4 stages. Yellow arrow shows PHB-AVA axonal de-fasciculation and corresponding exclusion of synapses. (**G, H**) Contact length (adjacency) between PHB and AVA neurons is reduced in *cdh-4* null mutants. Representative images (G) and quantification (H) of PHB-AVA CD4 GRASP reporter (*otEx8152*) in wild type and *cdh-4(ot1246)* null mutant animals at L1 and L4 stages. Dashed box represents the site of PHB and AVA contact. **(I)** ASK>AIA synapses are unaffected in *cdh-4(hd40)* mutants. **(J)** CRISPR-tagged CDH-4::GFP shows diffuse expression in the nerve ring. All images are maximum intensity projections of a subset of the Z-stack. Scale bars = 10μM. In all graphs, a dot represents one worm and error bars denote median with 95% confidence interval. P-values from unpaired t-test are shown.

To broaden our analysis to other neuron types, we assessed the role of *cdh-4/Fat* in forming synapses between additional neuron pairs. Synapses were unaffected in an ASK>AIA GRASP reporter in *cdh-4/Fat* mutants (**Figure 7I**). *cdh-4/Fat* null mutants have mispositioned IL2/IL1 somas - assessed by scoring the distance between IL2VR and IL1VR – with the distance between the two neuron somas being greater in mutants (**Figure S3D, E**). Despite the mispositioned somas, however, IL2>IL1 synapses - visualized with an NLG-1 GRASP reporter - could still be observed (**Figure S3F**).

Together with the lack of phenotype in PHB>AVA synapses, these results implicate CDH-4/Fat in mediating cell-specific neuronal soma positions and/or neurite contact, but not synapse specificity or synapse formation, at least in the specific contexts that we tested here. A role in determining neurite contact is also supported by CDH-4 localization patterns. A CRISPR-engineered CDH-4::GFP reporter shows visibly diffuse GFP expression along membranes in the nerve ring bundle at all stages of development (**Figure 7J**).

### The conserved cadherin *fmi-1/Celsr* controls axodendritic morphologies, axon pathfinding, and synapse formation

The broadly, neuronally expressed FMI-1/CELSR has several neuronal functions in *C. elegans*, as previously described, including axon navigation and synaptogenesis in the ventral nerve cord and sensory dendrite patterning (Steimel *et al*. 2010; Najarro *et al*. 2012; Hsu *et al*. 2020; Liao *et al*. 2024). To expand on the description of *fmi-1/Celsr* function beyond the nerve cord and sensory periphery, we further characterized *fmi-1/Celsr* null mutants in other parts of the nervous system. Using NeuroPAL, we found that neuronal soma positioning was largely unaffected in the head region in *fmi-1/Celsr* mutants (**Figure 8A**). In addition, overall nerve ring neighborhood topology was largely unperturbed based on AxoPAL analysis (**Figure 8B)**. However, a subset of neurons failed to enter the nerve ring (**Figure 8D-F**). For example, in a subset of animals, the AVE axon was missing from the anterior nerve ring neighborhood, which could be due to a defect in AVE neighborhood switch or its failure to enter the nerve ring. Similarly, inner labial neurons IL1/IL2 failed to enter the nerve ring in a subset of animals and their posteriorly projecting axonal loops were found much diverged from the posterior end of nerve ring (**Figure 8C**, worms #1 and #4). Hermaphrodite-specific midbody HSN neurons showed axon pathfinding defects as previously published (Steimel *et al*. 2010), and axons of male-specific EF neurons - whose cell bodies localize in the tail region and axons project to the nerve ring - failed to reach the nerve ring in a manner similar to the HSN phenotype (**Figure 8D, E**). AxoPAL and additional narrowly expressed cytoplasmic reporters revealed additional defects in axodendritic morphologies in *fmi-1/Celsr* mutants. For instance, in a subset of animals, IL1/IL2 neurons altogether lacked axonal loops that typically traverse the width of the nerve ring (**Figure 8C**, worm #5).

**Figure 8:**
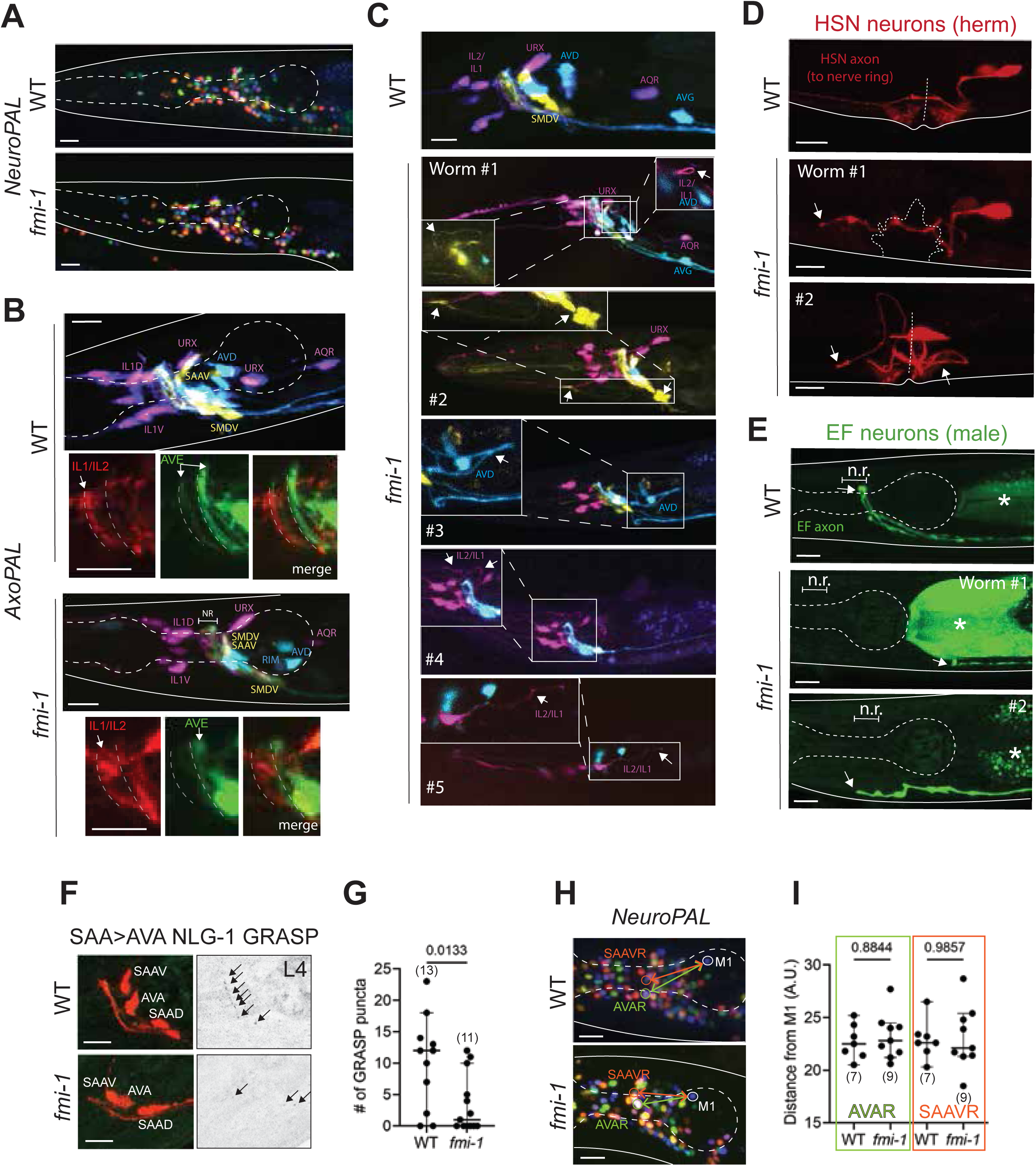
Conserved *fmi-1/Celsr* selectively mediates a subset of neuronal anatomical features. **(A)** Representative images of *NeuroPAL* reporter in the background of *fmi-1* null mutants. **(B)** Representative images of *AxoPAL* reporter in the background of *fmi-1(ot1090)* mutants. **(C)** Representative images of variable axodendritic defects in *fmi-1(ot1090)* mutants assessed with the *AxoPAL* reporter. **(D)** Representative images of variable axon pathfinding defects in hermaphrodite-specific HSN neurons in *fmi-1(ot1090)* mutants. **(E)** Representative images of variable axon pathfinding defects in male-specific EF neurons in *fmi-1(ot1090)* mutants. (**F, G**) SAA>AVA synapses are reduced in *fmi-1* null mutants. Representative images (G) and quantification (H) of SAA>AVA NLG-1 GRASP (*otIs839*) puncta in wild type and *cdh-4(ot1353)* null mutant animals at L1 and L4 stages. 10 animals per genotype were analyzed for the NeuroPAL and AxoPAL analyses. P-value from unpaired t-test is shown. (**H, I**) AVA and SAA soma positions are unaffected *fmi-1* null mutants. Representative images (G) and quantification (H) of shortest distance (yellow) between AVAR (green) and SAAVR (orange) soma labeled with *NeuroPAL* (*otIs669*) in wild type and *fmi-1* null mutant animals at the L4 stage. AVAR and SAAVR were scored relative to M1 pharyngeal neuron which does not express *fmi-1* and is unaffected in *fmi-1* mutants. P-values from unpaired t-test are shown.

We also assessed the role of *fmi-1/Celsr* in synapse formation. We have recently shown that *fmi-1/Celsr* is necessary and sufficient for establishing and maintaining neurite contact and proper synapses between neurons PHB and AVA (Liao *et al*. 2024). Here, we broadened our analysis of *fmi-1/Celsr’s* role in synapse formation as well as other aspects of neuronal architecture. We first used an NLG-1-based GRASP reporter that labels SAA>AVA synapses (Majeed *et al*. 2024), to assess how loss of *fmi-1/Celsr* affects other synapses. Indeed, we observed a significant reduction in SAA>AVA synapses in *fmi-1(ot1090)* null mutants (**Figure 8F, G**). However, SAA and AVA soma positions as well as their overall morphologies and placement in the nerve ring were unaffected by loss of *fmi-1/Celsr* (**Figure 8H, I**). It is possible that SAA>AVA synapses were lost as a consequence of loss of neurite contact, like in the case of PHB and AVA (Liao *et al*. 2024). These results, together with the *cdh-4/Fat* analyses described in the previous section, illustrate that different cadherins have distinct roles in nervous system development and that their effect on soma positions and neurite positions can be decoupled from proper synapse formation in the paradigms studied.

### Mapping cadherin expression onto the synaptic connectome

We next turned from the broadly expressed and conserved cadherins *cdh-4/Fat*, and *fmi-1/Celsr,* to the remaining neuronal cadherins that are all much more restrictively expressed, namely, the conserved cadherins *cdh-1/Dchs* and *cdh-3/Fat,* and the *C. elegans*-specific *cdh-5, cdh-8, cdh-9* and *cdh-12* genes (**Figure 2A-C**). We first mapped their nervous system-wide expression onto the *C. elegans* synaptic connectome of the adult (**Figure 9A**)(Cook *et al*. 2019). This mapping is particularly suitable to visualize several points already mentioned above (**Figure 9A)**: (a) in aggregate, neuron-specific cadherins do not cover the entire nervous system; (b) there is a finite number of neurons that express multiple cadherins and (c) cadherin expression is not biased toward specific parts of the nervous system or does not obviously correlate with specific connectivity features of a neuron. Most importantly, though, such mapping visualizes whether neurons that express a given cadherin are synaptically connected to neurons that express the same cadherin. Due to the well-documented ability of cadherins to engage in homophilic interactions (Shapiro *et al*. 1995a; Brasch *et al*. 2018; Honig and Shapiro 2020), such matching expression may indicate a role in synapse formation or function.

**Figure 9:**
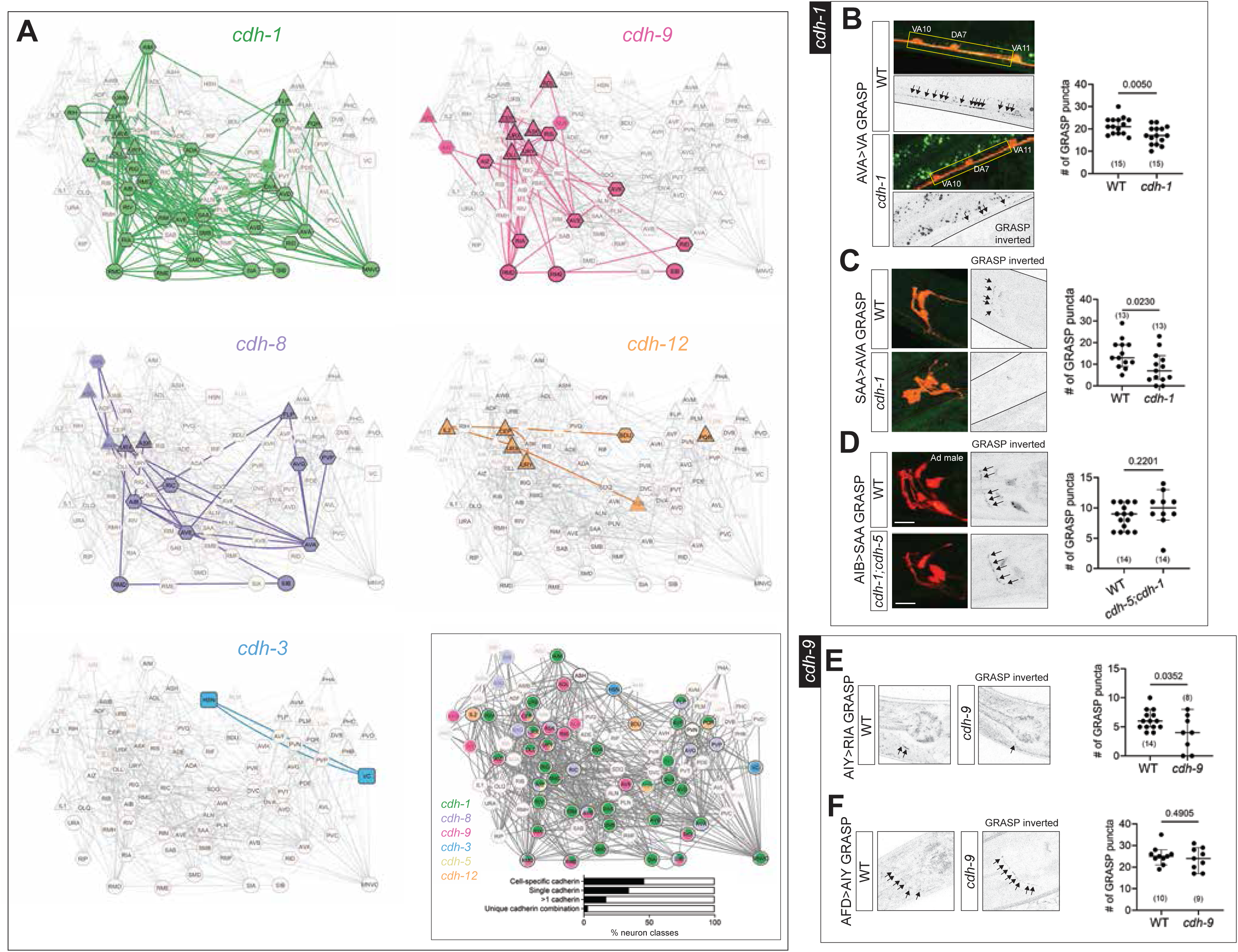
Neurons expressing narrowly expressed cadherins are synaptically connected and mediate specific synapses. **(A)** Expression of *cdh-1*, *cdh-9*, *cdh-8*, *cdh-12*, and *cdh-3* overlaid on the adult connectome (Cook *et al*. 2019). MNVC includes all ventral nerve cord motor neurons except VC. Boxed panel shows expression of all uniquely and narrowly expressed cadherins overlaid on the adult connectome. Percentages of neuron classes expressing cell-specific cadherins, single or multiple cadherins, and unique cadherin combinations are shown. Triangle, square, and hexagon nodes represent sensory, motor, and interneurons. Edges represent synaptic connectivity (unweighted) based on Cook et al., 2019. **(B)** AVA>motor neuron (VA) synapses are reduced in *cdh-1(ot1034)* null mutants. Representative images and quantification in L4 stage animals is shown. Yellow box represents the site of synapses. **(C)** SAA>AVA synapses are reduced in *cdh-1(ot1034)* null mutants. Representative images and quantification in L4 stage animals is shown. **(D)** AIB>SAA synapses are unaffected in *cdh-1(ot1034); cdh-5(ot1093)* double mutants. Representative images and quantification in adult stage males is shown. **(E)** AFD>RIA synapses are reduced in *cdh-9(ot1247)* null mutants. Representative images and quantification in L4 stage animals is shown. **(F)** AFD>AIY synapses are unaffected in *cdh-9(ot1247)* null mutants. Representative images and quantification in L4 stage animals is shown. All images are maximum intensity projections of a subset of the Z-stack. Scale bars = 10μM. In all graphs, a dot represents one worm and error bars denote median with 95% confidence interval. P-values from unpaired t-test are shown.

With the obvious exception of *cdh-5,* which is only expressed in a single neuron class, it is indeed apparent that almost all the neuron type-specifically expressed cadherins are expressed in neurons that are synaptically connected to neurons that express the same cadherin (**Figure 9A**). To take animal to animal variability of synaptic connectivity into account (Witvliet *et al*. 2021), we overlaid cadherin expression on a wiring diagram that only contains synaptic connections that are present in all available *C. elegans* datasets of larval-stage and adult animals, of either sex (Cook et al., 2023). In such a “core connectome”, the theme of cadherin-expressing neurons being synaptically connected to neurons expressing the same cadherin still largely holds (**Figure S4**).

### Loss of the conserved cadherin *cdh-1/Dchs* and *C. elegans*-specific cadherin *cdh-9* selectively result in synaptic defects

Together with their predicted ability to engage in homophilic protein interactions, the expression of individual cadherins in neurons that are directly connected to one another provide testable hypotheses for potential roles of cadherins in controlling synaptic connectivity and/or synaptic function. We considered cadherins (*cdh-1/Dchs, cdh-9, cdh-3/Fat*) for experimental analysis, using previously described GRASP-based markers for synaptic connectivity. Using NeuroPAL and AxoPAL (for *cdh-1/Dchs* and *cdh-9)* or neuron-specific markers (for *cdh-3/Fat),* we examined neuronal soma position and neurite neighborhood/projections of neurons expressing *cdh-1/Dchs, cdh-9* or *cdh-3/Fat* in the absence of the respective genes and found no obvious defects (**Figure S5, S6A)**.

Based on GRASP marker availability, we examined connectivity of the synaptically connected neuron classes SAA, AVA, and VA, which all express *cdh-1/Dchs* (**Figure 9A**). We found a reduction of synaptic puncta in AVA>VA and SAA>AVA synapses trans-synaptically marked with GRASP (**Figure 9B, C)**. Synaptic contacts made by SAA to another neuron, AIB, are not affected (**Figure 9D)**.

In the case of *cdh-9*, we examined its role in specifying the synapses between the interconnected, *cdh-9-*positive neurons AFD, AIY and RIA. GRASP reagents are available to visualize AFD>AIY and AIY>RIA synapses (Feinberg *et al*. 2008). In the case of AIY>RIA synapses located in the ventral region of the nerve ring, we observed a reduction in connectivity in *cdh-9* mutants; puncta were fewer and dimmer in mutants compared to wild type animals (**Figure 9E)**. AFD>AIY synapses were unaffected in *cdh-9* mutants (**Figure 9F)**, implicating *cdh-9* as a synaptic effector of neuron-specific synapses, like in the case *cdh-1*. Notably, an absence of synaptic phenotype in the AFD>AIY reporter suggests that the overall axonal morphology of AIY neurons is unperturbed in *cdh-9* mutants.

Lastly, we considered the *cdh-3/Fat* cadherin, which within the nervous system is exclusively expressed in the synaptically interconnected HSN and VC neurons that form the circuit for egg-laying behavior (**Figure 2A, B**) (Schafer 2005). Bright HSN and VC expression of *cdh-3/Fat* was detected at the L4/Ad stage, when the egg-laying circuit matures. HSN synapses onto vulval muscles (Vm) and VC neurons in the vulval region. We assessed HSN>Vm and HSN>VC synapses, labeled previously with trans-synaptic GRASP reporters, but observed no defects in the number or distribution of synaptic punctae (**Figure S6A, B**). A marker of presynaptic active zones, CLA-1::GFP, specifically expressed in HSN also showed no defects (**Figure S6C**); CLA-1 puncta were present and localized properly in the vulval region, in contrast to the ectopic presynaptic specializations observed in other cell adhesion proteins mutants (Shen and Bargmann 2003). In addition to vulval synapses, HSN forms synapses with several neuron classes in the nerve ring region (White *et al*. 1986). While we did not test a role of *cdh-3/Fat* in specific nerve ring synapses using GRASP-based reporters, CLA-1::GFP puncta in the nerve ring were unaffected in *cdh-3(ot1035)* mutants (**Figure S6D**). Since we find *cdh-3/Fat* null mutant to display an egg retention defect akin to (but less severe) than HSN loss (**Figure S6E**), it is possible that similar to the function of other cadherins, *cdh-3/Fat* may affect synaptic transmission rather than synaptic morphology of the egg laying circuit. We caution that the expression of *cdh-3/Fat* in other cells of the egg-laying system (**Figure 3B**) (Pettitt *et al*. 1996) may provide an alternative explanation of the egg retention defects of *cdh-3/Fat* mutants.

### *C. elegans*-specific cadherin CDH-5 is required for proper electric synapse patterning of the AIB interneurons

We discovered a synaptic role for another cadherin, the nematode-specific *cdh-5* gene, which is the only cadherin that is neuronally expressed exclusively in a single pair of neurons, the interneuron pair AIB (**Figure 2A, S2B**). The neurites of AIB neurons project into the nerve ring, where they generate both chemical and electrical synaptic connections (White *et al*. 1986). Food scarcity and crowded external conditions, prompting animals to enter the dauer diapause state, trigger AIB to generate additional electrical synapses (Bhattacharya *et al*. 2019). Cadherins have been localized to electrical synapses in the vertebrate nervous system (Cardenas-GARCIA *et al*. 2023), but it is not known whether cadherins are required for assembling neuronal electrical synapses.

Examining *cdh-5* reporter allele expression in dauer stage animal, we found that *cdh-5* is upregulated in AIB neurons in dauers (**Figure 10A, B)**. This upregulation correlates with the dauer-specific induction of the electrical synapse-forming INX-6 innexin exclusively in the AIB neurons (Bhattacharya *et al*. 2019). Assessment of AIB-localized INX-6::GFP puncta in *cdh-5(ot1093)* null mutants revealed a significant reduction in puncta in mutant dauers relative to wild type control dauers (**Figure 10C, D**). Restoring CDH-5 specifically in AIB interneurons using an AIB-specific promoter (*inx-1p*) to drive full-length CDH-5 genomic DNA rescued the phenotype of *cdh-5* mutant dauers in two independent transgenic lines (**Figure 10C, D**).

**Figure 10:**
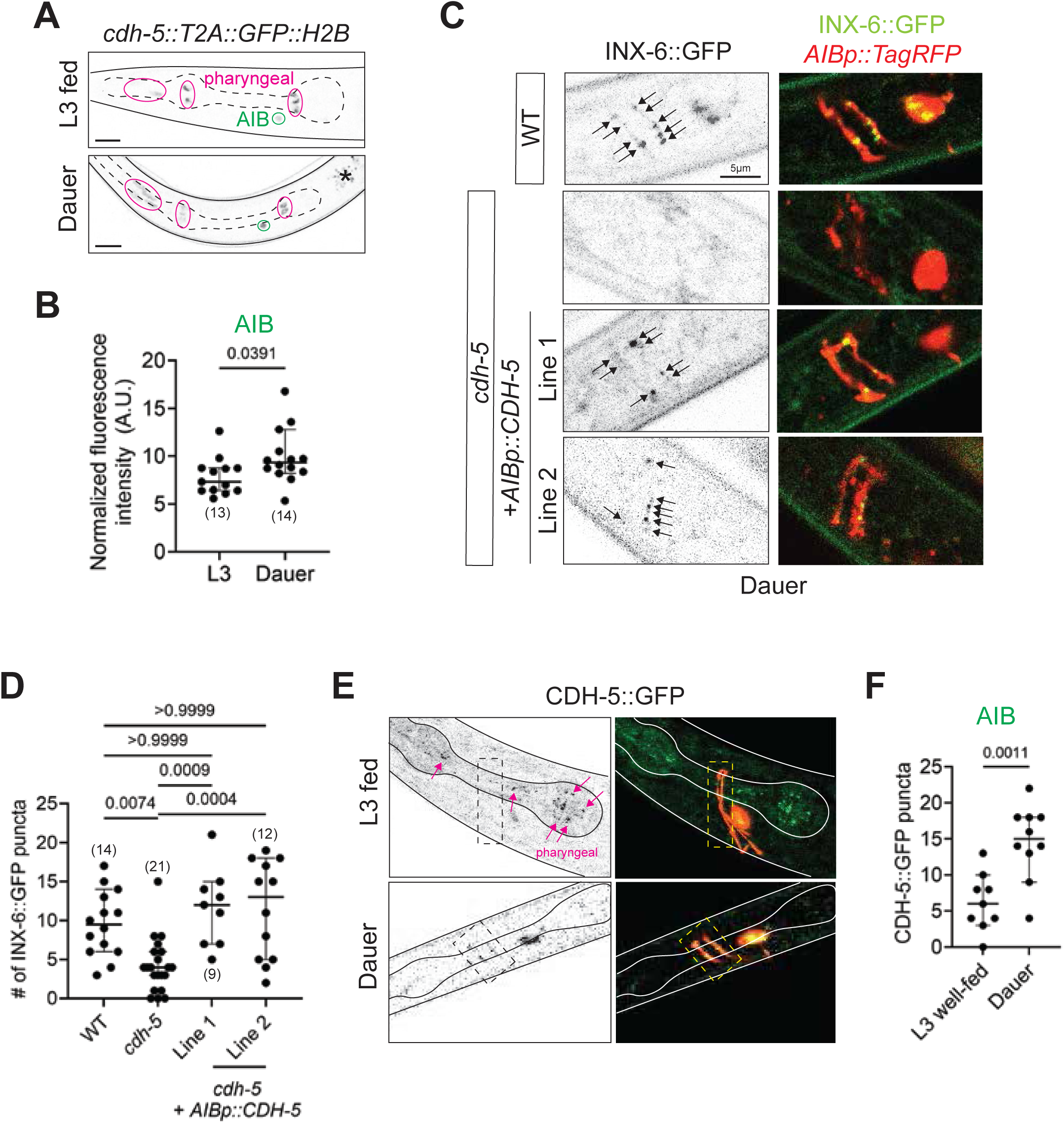
*C. elegans*-specific cadherin *cdh-5* is required for electrical synapse formation in dauers. **(A, B)** Experience-dependent plasticity of *cdh-5* expression in AIB neurons. Representative images (A) and quantification (B) of *cdh-5*(*ot1127*) expression in AIB interneurons shows enrichment at the stress-induced dauer stage relative to well-fed L3-stage control. Images denote AIB and non-neuronal *cdh-5* expression in green and pink, respectively. Normalized fluorescence intensity in the AIB neuron closer to the microscope lens in lateral worms is quantified. P-values from unpaired t-test are shown. (**C, D**) Representative images (C) and quantification (D) of INX-6::GFP (*ot805*) puncta in wild type and *cdh-5(ot1316)* null mutant dauer worms. Complete loss of CDH-5 leads to a decrease in INX-6::GFP puncta that colocalize with AIB axons. AIB-specific expression of CDH-5 cDNA using *inx-1prom* (*otEx8156*) rescues the loss of INX-6::GFP phenotype in 2 independent transgenic lines. AIB neurons were cytoplasmically labeled with *npr-9p::TagRFP* (*otEx8072*) and only colocalizing INX-6::GFP puncta were scored. P-values shown were calculated using one-way ANOVA with Bonferroni correction for multiple comparisons. Scale bar = 5μM. (**E, F**) Representative images (E) and quantification (F) of CDH-5::GFP (*syb6640*) puncta localizing on AIB axons labeled with *npr-9p::TagRFP*. Number of CDH-5::GFP puncta increase in dauers relative to L3 well-feed control animals. P-values from unpaired t-test are shown. All images are maximum intensity projections of a subset of the Z-stack. Scale bars = 10μM unless otherwise noted. In all graphs, a dot represents one worm and error bars denote median with 95% confidence interval.

We generated a CRISPR-engineered CDH-5::GFP reporter in which *gfp* is inserted between the PDZ motif-containing C-terminus and transmembrane domain to 1) tag all isoforms and 2) avoid disrupting endogenous protein localization. We observed punctate CDH-5 expression along AIB axons (as well as the pharynx, where INX-6 is also expressed; (Bhattacharya *et al*. 2019)). In line with dauer-specific upregulation of our nuclear *cdh-5* reporter, we found an increase in CDH-5::GFP puncta localized on AIB axons in dauer animals (**Figure 10E, F**). While the similar fluorophores of the CDH-5 and INX-6 reporter alleles prevented the assessment of an overlap of these two factors, the punctate localization of both proteins is consistent with a synaptic localization pattern.

Aside from electrical synapses, we characterized *cdh-5* mutants for defects in AIB soma positions, axon topology in the nerve ring, AIB-specific presynaptic specializations marked with CLA-1::GFP, and specific synapses between AIB and a subset of its foremost chemical synaptic partners (RIM and SAA) labeled with iBLINC and GRASP, respectively, and found no significant phenotypes (**Figure S7**). These observations indicate that *cdh-5* does not disrupt outgrowth, neighborhood or chemical synaptic target choice of the AIB axon but apparently has a selective role in promoting electrical synapse formation.

## DISCUSSION

We have provided here a broad, genome- and nervous system wide view of the expression and function of the cadherin gene family in *C. elegans.* This broad view reveals several themes that we first discuss from the perspective of gene expression patterns and then from the perspective of gene function.

### Dichotomies of cadherin gene expression patterns in the nervous system

Our expression pattern analysis of the entire cadherin gene family, conducted at several stages of development has revealed several themes.

First, there are two dichotomies in the expression pattern of the 12 *C. elegans* cadherin genes: (a) only two are expressed during proliferative and migratory stages of embryogenesis (classic cadherin *hmr-1* and *cdh-4/Fat*); the expression of all others is only initiated about the time when cells exit the cell cycle and morphogenesis begins, (b) within the nervous system, 4 of the 12 genes are expressed in an essentially panneuronal manner, while 6 cadherins show a very restricted expression in the nervous system (the remaining 2 are not expressed at all in the nervous system).

Second, the dichotomy of panneuronal and very selectively expressed cadherins almost perfectly correlates with the conservation of the cadherins: All 4 panneuronal cadherins are conserved, while all the non-conserved cadherins are very narrowly expressed in the nervous system. There are only two neuron-selectively expressed cadherins that are phylogenetically conserved (*cdh-1/Dchs* and *cdh-3/Fat*).

Third, only a limited fraction of neurons (46%) expresses a neuron-type specific cadherin and expression of the neuron type-specific cadherins show only limited overlap. Hence, there are few neurons that express unique combinatorial codes of neuron type-specific cadherins.

Fourth, while there are some temporal dynamics of onset of cadherin expression in the embryo, as well as several changes during larval development, all neuronally expressed cadherins show expression in the fully mature, adult nervous system, lasting throughout the life of the animal.

These expression patterns make several predictions about the spectrum of cadherin gene function. First, the broadly expressed cadherins may be involved in “generic”, i.e. non cell-type specific aspects of cellular morphogenesis or behavior. These generic functions may be implemented in a cell-type specific manner through the interaction of a broad cadherin with cell-type specific partner proteins. Second, expression in the adult nervous system indicates postdevelopmental roles and, third, since there are very few cell-specific combinations of cadherins, it is unlikely that cell-specific cadherin codes define neuron-type specific connectivity features on a broad scale, as they do, for example, in the vertebrate retina (Duan *et al*. 2018; Sanes and Zipursky 2020). Together with the notion that many cells do not even express any cell-type specific cadherin, there is clearly no cadherin-based logic that can comprehensively explain synaptic connectivity throughout the entire nervous system of *C. elegans*. While this could be viewed as an overly naïve hypothesis to begin with, one should be reminded that cadherins predate the advent of nervous system and even multicellularity. It therefore appears reasonable to envision that through the expansion of this gene family from a primitive ancestor, cadherin deployment became a pervasively used mechanism for increasing the complexity of cellular interactions in a nervous system. However, at least in *C. elegans*, neuron-specific cadherins only cover parts of the nervous system.

### Cadherin functions on several anatomical scales

Our gene family-wide mutant analysis reveals that the cadherin expression dichotomy correlates with function. Only two of the *C. elegans* cadherin genes, cadherin *hmr-1* and *cdh-4/Fat,* both phylogenetically deeply conserved, are essential for viability. These happen to be the only cadherins that are already expressed in proliferative embryonic stages during cell focusing and gastrulation. The phenotypes of all other cadherins, including the two panneuronal and conserved cadherins *fmi-1/Celsr* and *casy-1/Clstn* are strikingly subtle on a whole animal level. All animals appear morphologically normal and do not display any gross abnormalities normally associated with strong defects in nervous system development and function, such as the uncoordinated locomotion defects observed upon defects in synaptic transmission and/or failure of the motor system to properly develop. Behavioral phenotypes for the two panneuronal, conserved cadherins *fmi-1/Celsr* and *casy-1/Clstn* only become apparent upon assessing more sophisticated levels of nervous system function, such as complex associative learning paradigms (Ikeda *et al*. 2008; Hoerndli *et al*. 2009; Thapliyal *et al*. 2018a; Thapliyal *et al*. 2018b).

Despite the absence of gross behavioral defects, a more detailed anatomical analysis of the consequences of cadherin removal provides a nuanced and panoramic view of cadherin function in *C. elegans,* illustrating that they act on multiple organizational levels and anatomical scales of nervous system development. Some cadherins appear to function only on some anatomical scales, while others operate on all organizational levels. The latter refers explicitly to the phylogenetically conserved, and panneuronally expressed *cdh-4/Fat* gene. Previous analysis of *cdh-4/Fat* mutants (Schmitz *et al*. 2008; Sundararajan *et al*. 2014), expanded here by our analysis of *cdh-4/Fat* throughout different parts of the nervous system, using multiple cutting-edge imaging tools and approaches, reveals that *cdh-4/Fat* is required for proper placement of neuronal soma, for proper outgrowth, placement and relative positioning of neurites within fascicles and for synaptogenesis. Intriguingly, at least some of these functions appear to be executed separately from each other, such that defects in synapse formation are not necessarily a consequence of earlier outgrowth defects and vice versa, that defects in soma positioning do not necessarily lead to defects in synaptogenesis. The phenotypic spectrum of another broadly expressed and conserved cadherin, *fmi-1/Celsr* is somewhat narrower (i.e. animals are viable and show very limited overall tissue organization defects), but *fmi-1/Celsr* still affects several levels of circuit formation, from axon outgrowth to synapse formation. Such a function of *fmi-1/Celsr* has also recently been delineated in the context of sexual dimorphic maturation of the nervous system (Liao *et al*. 2024).

The more narrowly expressed cadherins show phenotypes that are limited to specific organizational levels. Perhaps most interestingly, we observed synaptogenic functions of several conserved and non-conserved cadherins that are accompanied by no obvious defects in other aspects of neuronal morphogenesis. These defects occur in cellular contexts where the respective cadherin is expressed on both the pre- and postsynaptic side of connected cells, implying a homophilic function for these cadherins. Whether these synaptic defects reflect Sperry-type lock-and-key synaptic recognition events or the result of cadherins involved in determining the extent of neurite neighborhood (Cook *et al*. 2023; Wolterhoff and Hiesinger 2024) remains to be determined. We note that the effects of cadherins on synaptic read-outs are not fully penetrant, indicating that other factors likely play an important role, too.

While functions of cadherins in chemical synapse formation (Williams *et al*. 2011; Kuwako *et al*. 2014; Basu *et al*. 2017; Duan *et al*. 2018; Yamagata *et al*. 2018; Zou 2020) are surely not unprecedented, we discovered here a novel and unprecedented role of a cadherin, *cdh-5*, in electrical synapse formation. It is conceivable that the function for cadherins in localizing innexins may extend to other electrical synapses as well. But, as much as we argue (based on expression patterns) that cadherins are unlikely to play a pervasive role in the formation of all chemical synapses, a broad and invariant role of cadherins in electrical synapse formation appears unlikely as well, again based on cadherin expression patterns, but also based on the notion that the phenotypic spectrum of innexin mutant animals (Jin *et al*. 2020) is much broader than that of cadherin mutant animals.

### Functions of cadherins in the mature nervous system

One of the most striking aspects of the expression of any *C. elegans* cadherin is their maintained expression in the mature nervous system throughout the life the animal. Contrasting their well-mapped expression during *Drosophila* and vertebrate embryonic development, cadherin family expression in the mature nervous system has not been comprehensively explored in other organisms. In those cadherin mutants in which we have not observed any obvious developmental or morphological defects, such as in *cdh-3/Fat* mutants, we infer that cadherins may dictate selective aspects of neuron function that is required throughout the life of the neuron. One precedent for such a scenario is the *C. elegans* calsyntenin gene *casy-1/Clstn*, which in specific cellular settings is required for synaptic transmission and plasticity (Ikeda *et al*. 2008; Hoerndli *et al*. 2009; Ohno *et al*. 2014; Thapliyal *et al*. 2018a; Thapliyal *et al*. 2018b). Functions of several vertebrate cadherins in synaptic signaling, homeostasis and maintenance have been described (Friedman *et al*. 2015) and we consider those to be the most likely functions of the *C. elegans* cadherins for which we have not found a function yet.

### Similarities and dissimilarities between the function of animal cadherins

There are striking parallels of *C. elegans* and vertebrate cadherin function, but besides pointing to novel aspects of cadherin function (e.g. in electric synapse formation), our analysis also reveals striking differences. First and foremost, on the level of existence of genes, there are no protocadherins in *C. elegans*. In vertebrates, protocadherins are thought to be involved in self-avoidance of sister neurites of a developing neuron (Zipursky and Sanes 2010; Mountoufaris *et al*. 2018). The much less elaborated nature of *C. elegans* neurites (White *et al*. 1986) is a possible explanation for why protocadherins do not exist in *C. elegans*.

Several of the cadherins conserved between worms and vertebrates have similar functions. For example, there are common roles of FMI-1 and its vertebrate homolog CELSR in axon pathfinding and synaptogenesis (Steimel *et al*. 2010; Najarro *et al*. 2012; Chai *et al*. 2014; Thakar *et al*. 2017; Liao *et al*. 2024) and CDH-4 and its vertebrate homolog FAT in neuronal migration and axon pathfinding (Schmitz *et al*. 2008; Sundararajan *et al*. 2014; Aviles and Goodrich 2017; Fulford and Mcneill 2020). On the other hand, there are also apparent functional discrepancies that may point to novel and currently unexplored aspects of vertebrate cadherin function. For example, in both *Drosophila* and vertebrates, FAT-type cadherins operate together with a DCHS/Dachsous-type cadherin as a ligand/receptor pair to fulfill a number of diverse roles during morphogenesis and embryonic pattern formation, within and outside the nervous system (Aviles and Goodrich 2017; Fulford and Mcneill 2020). Yet in *C. elegans*, we found that loss of the sole Dachsous ortholog, *cdh-1* (which had not been studied before), results in none of the obvious neuronal developmental and morphological defects of *cdh-4/Fat* mutants. Moreover, unlike *cdh-4/Fat* mutants, *cdh-1/Dchs* mutants are completely viable. This lack of matching phenotype is also consistent with the strikingly more restricted expression of *cdh-1/Dchs.* These observations strongly argue for FAT proteins having DCHS-independent function, possibly as homophilic adhesion molecules or via binding partners that are unexplored in any animal system (Aviles and Goodrich 2017).

In conclusion, by taking a panoramic genome- and animal-wide approach, we identify roles of cadherins in novel cellular contexts. We find that cadherin functions at various anatomical scales (animal morphogenesis, behavior, soma positions, axon patterning, synapses) to orchestrate the development of some components of the *C. elegans* nervous system. Our comprehensive expression atlas provides an entry point for exploring cadherin function more deeply and in additional contexts, such as in non-neuronal tissues, during synapse maintenance and experience-dependent behavioral plasticity, or during sexually dimorphic nervous system maturation.

## ACKNOWLEDGEMENTS

We thank Chi Chen for assistance with microinjections to generate strains and Steven J. Cook for valuable input on constructing connectivity maps. We thank Chun-Liang Pan, Hannes <u>Bülow</u>, Harald Hutter and members of the Hobert lab for comments on the manuscript. This work was funded by the HHMI and by the Charles Revson postdoctoral fellowship for CPL.

## MATERIALS AND METHODS

### *Caenorhabditis elegans* strains and maintenance

Worms were grown at 20°C on nematode growth media (NGM) plates seeded with *E. coli* (OP50) bacteria as a food source, as previously described (Brenner 1974). Worms were maintained according to standard protocol. The wild-type strain used in this study denotes *C. elegans* Bristol variety (N2). A complete list of strains used in this study is listed **Table S2**. Strains generated in this study are being deposited at the CGC.

### CRISPR/Cas9-based genome engineering

To generate endogenously tagged T2A::GFP::H2B cadherin reporters as previously described (Ghanta *et al*. 2021), the following sgRNAs were used:

<u>*cdh-1*:</u> 5’AACGATGGAATGATTGGTGG3’

<u>*cdh-3*:</u> 5’GCCCCATCTTACCGTAGAGA3’

<u>*cdh-5*:</u> 5’CGTTGATGATAATCTGGTGT3’

<u>*cdh-8*:</u> 5’GACTGCAAATCTGCAGAAGT3’

<u>*cdh-9*:</u> 5’acataTTAAAAATAGACAGT3’

<u>*cdh-12*:</u> 5’CATCGTTGGCAATTGATGGG3’

<u>*casy-1*:</u> 5’AGAACGAGCGTTCGTTGAGA3’ and 5’GCAAATCAACGTGTCGTTGG3’

To generate full locus cadherin deletion alleles, two guide sgRNAs at either end of the locus were used in a single injection as previously described (Bayer and Hobert 2018), as follows:

<u>*cdh-1(ot1034* and *ot1299)*:</u> 5’ TTGGGAACATGATGTTTCGG 3’ and 5’ AACGATGGAATGATTGGTGG 3’

<u>*cdh-3(ot1102* and *ot1248)*:</u> 5’CGAAAACTTGATCGTCTCTT3’ and 5’GCCCCATCTTACCGTAGAGA3’

<u>*cdh-3(ot1035)*:</u> 5’ CGAAAACTTGATCGTCTCTT3’ and 5’ CTTCTATGAAAATAGTTGCA3’

<u>*cdh-4(ot1246* and *ot1353)*:</u> 5’AGTCACCACATCTTCTTCTA3’ and 5’GTCTGTGAATCTAATGAGAT3’

<u>*cdh-5(ot1093)*:</u> 5’TGTTTCAGTTAAGAACCCTT3’ and 5’CGTTGATGATAATCTGGTGT3’

<u>*fmi-1(ot1090* and ):</u> 5’TTGAATGTGAATGTCAGTGG3’ and 5’TGATGCGTATTACACATATA3’

<u>*cdh-8(ot1084* and *ot1319)*:</u> 5’GGTGTCTAAAAGGAACAGGT3’ and 5’GACTGCAAATCTGCAGAAGT3’

<u>*cdh-9(ot1247* and *ot1300)*:</u> 5’ TAGTAAGGATCGGCCCAAGT 3’ and 5’ACATATTAAAAATAGACAGT3’

<u>*cdh-12(ot1091)*:</u> 5’CTGCGAATAATAGTATGAAG3’ and 5’CATCGTTGGCAATTGATGGG3’

<u>*casy-1(ot1082)*:</u> 5’AGCATGGTGATGTTTGGCGT3’ and 5’GCAAATCAACGTGTCGTTGG3’

To generate *unc-86* deletions *ot1355* and *ot1354* in the background of *cdh-12(ot1119)* and *fmi-1(syb4563)* GFP reporters, respectively, the following sgRNAs and ssODN were used:

sgRNA 1: 5’CAAGGTCCCCCTCTTTTCCA3’

sgRNA 2: 5’ACAACATACAATGGGCTACC3’

Repair template: 5’TCTGTCTCCTCCCAGCTTCAAGGTCCCCCTCTTTTACCTTGATTCTTTGATTAGTTTCGTTTTCGTGAAC3’

CRISPR-tagged SL2::GFP::H2B reporters for *fmi-1(syb4563)*, *hmr-1(syb4454)*, *cdh-4(syb4454)*, and *cdh-7(syb4675)* were made by SunyBiotech. The CDH-4::GFP CRISPR reporter (*syb6764*) was also made by SunyBiotech.

### Molecular cloning

#### AxoPAL

Transgenic reporter *otEx7895* (AxoPAL v1.0) contains the following reporters: nmr-1::CyoFP (AVA, AVE, AVD, RIM, AVG, PVC), flp-3::mNeptune (IL1), gcy-35::mNeptune (URX, AQR, ALN, PQR, PLN, SDQ), flp-7::tagRFP (SAA), klp-6::tagRFP (IL2), and C42D4.1::GFP (RMD, SAA). Promotors were either cloned using standard restriction digestion cloning or excising NLS/H2B sequence from plasmids used to generate NeuroPAL (Yemini et al., 2021). All plasmids were co-injected in N2 strain at 30ng/µL each; no coinjection marker was used because the cytoplasmic reporter expression was visibly bright. Trasgenic worms were maintained by picking fluorescent worms.

Primers for respective gene promoters used in AxoPAL v1.0:

*nmr-1p* – Forward: gaaatgaaataAGCTTGCATGCCTGCAGctgctgctgtaggctttg, Reverse: GTCCTTTGGCCAATCCCGGGatctgtaacaaaactaaagtttgtcg; 2xNLS, H2B were excised from pEY32 (Addgene # 191037, (Yemini *et al*. 2021)) using Gibson.

*gcy-35p* - Forward: gaaatgaaataAGCTTGCATGCCTGCagtttccgcatatagcttatag, Reverse: GTCCTTTGGCCAATCCCGGGattctactctccgcaaaaaagtaac; 3xNLS was excised from plasmid pEY45 (Addgene # 191049, (Yemini *et al*. 2021)) using Gibson.

*klp-6p* (pCC045) - Forward: cgttttggagtttgctacga, Reverse: tattctgaaaagttcaactaataaatttag

*flp-3p* - Forward: gaaatgaaataAGCTTGCATGCCTGCAGACTAgttccgcatggaataatagcc, Reverse: GTCCTTTGGCCAATCCCGGGtggtggttatggtggtgttac. 3xNLS was excised from pEY44 (Addgene # 191048, (Yemini *et al*. 2021)) using Gibson.

*flp-7p* – Forward: actgttgcttggtcttgtg, Reverse: ttctaaaagtctttgaatgaaaacgag *C42D4.1p* (pCC277, (Cros and Hobert 2022))

#### *cdh-5* rescue

Genomic *cdh-5* sequence was amplified from wild type N2 lysate and cloned into *inx1p::SL2::TagRFP* plasmid using Gibson cloning (NE Builder HiFi DNA Assembly Master Mix, E2621L).

### Expression analysis and cell identification

Cell-specific neuron identification for all cadherin reporters was performed by colocalization with the NeuroPAL landmark reporter transgene (*otIs669*) as described previously (Yemini *et al*. 2021). Briefly, each cadherin reporter strain was crossed into NeuroPAL strain, and 10-15 animals were imaged for analysis. Animals were imaged in the lateral position to facilitate identification. For L1 scoring, some animals were imaged in a dorsoventral position to facilitate neuronal identification in crowded ganglia. Synchronized L1- and L4-stage animals were obtained by egg prep.

For non-neuronal cell identification, cadherin-positive cells were identified by assessing a combination of features including position, subcellular features based on Nomarski optics (Yochem, J 2006 *WormBook*; Mango, SE 2007 *WormBook*; (Pham *et al*. 2021), and non-overlap of GFP with the pan-neuronal TagRFP transgene in NeuroPAL. In both neuronal and non-neuronal cells, expression was scored as present or absent, irrespective of strength of fluorescent signal, if it was consistent across animals 5-10 animals.

### Automated worm tracking

#### Crawling assay

Crawling locomotion features were analyzed using an automated multiworm tracker system (MBF BioScience) and WormLab software (MBF BioScience). Briefly, 5-8 L4 hermaphrodite animals were placed on 5 cm NGM plates and videos were recorded for 4min at room temperature. Videos were captured at 7.7μm/pixel camera resolution to largely cover a 5cm plate (ROI). NGM plates were fully coated with a thin lawn of OP50 16-18h prior to the assay to prevent bacterial overgrowth and avoid biased locomotion at the edge of the bacterial lawn. L4 animals were synchronized via egg prep, and well-fed animals were used for tracking. Mutant and wild type N2 control animals were tracked on the same day by alternating between the two genotypes. Altogether, X animals were used for each condition.

#### Swimming assay

Swimming assay was performed as previously described using the WormLab automated multi-worm tracker (MBF BioScience) (Restif *et al*. 2014). Briefly, 5 L4 hermaphrodite animals, synchronized via egg prep, were transferred to 50 M9 buffer and recorded for 1min. Quantitative analysis of swimming locomotion was performed using the WormLab software (MBF BioScience). Worms that did not move right after placing in M9 were discarded from the further analysis. Animals from multiple plates were pooled and population-level swimming features - including speed, wave initiation rate, activity index, and curling - as described previously (Restif *et al*. 2014), were analyzed for statistical significance.

### Dauer selection

For *cdh-5* analyses, dauers were induced under starvation, crowding, and high temperature conditions. Briefly, 10 L4 hermaphrodite animals were placed on regular 5 cm NGM plates seeded with *E. coli* (OP50) and incubated at 25°C. 1.5 weeks later, the plates contained a mixed population dauers and non-dauers. To isolate enough dauers for analysis, sodium dodecyl sulfate (SDS)-selection was performed as previously described (Cassada and Russell 1975). Briefly, a mixed population of animals was washed off plates, centrifuged, resuspended in 1% SDS, and incubated at room temperature with gentle agitation. 30min later, dauer larvae were centrifuged, washed twice with M9 to remove SDS, and transferred to uncoated NGM plates. All experiments were performed within 1 hour of transfer to prevent animals from exiting dauer arrest. To compare dauer animals to L3 well-fed wild type N2 control, L3 animals were synchronized via egg prep.

### Microscopy

Worms were anesthetized using 100mM sodium azide (NaN) and mounted on glass slides with 5% agarose pads. Images were acquired using a confocal laser scanning microscope (Zeiss LSM880) and analyzed using ZEN Imaging software (blue edition, Carl Zeiss) or Fiji ImageJ (Schindelin *et al*. 2012). NeuroPAL and AxoPAL images were acquired at 40x. Synaptic reporter images were typically acquired at 63x.

### Statistical analysis

All statistical analysis was performed using Prism (GraphPad). Details of analyses are mentioned in figure legends. For tracking analyses, locomotion metrics were first exported from WormLab to Excel, and statistical tests were subsequently performed in Prism.

### Analysis of AxoPAL neighborhood topology

Axonal topology was visualized in Fiji by first cross-sectionally slicing the region of the nerve ring and max-projecting the resliced region onto the Z-plane in increments spanning the nerve ring width to assess nerve ring compoisiton along the anterior-posterior axis.

### Quantification of synaptic puncta

ASK>AIA, PHB>AVA, BDU>HSN, SAA>AVA, and AIB>SAA GRASP puncta were scored using the semi-automated worm puncta scoring software WormPsyQi (Majeed *et al*. 2024). In all cases, a neurite mask was not used for synapse segmentation except in the case of AIB and NSM CLA-1 datasets.

AIB>RIM iBLINC puncta were scored by first cross-sectionally slicing the region of the nerve ring containing iBLINC puncta, making it in the Z-plane, and then counting manually in Fiji. AFD>AIY, AIY>RIA, HSN>Vm, and HSN>VC were scored using the cell counter plugin in Fiji.

CLA-1 puncta were scored for overall number and puncta features - such as mean pixel intensity and mean inter-synapse distance - using WormPsyQi.

### Quantification of PHB and AVA contact

Contact site length between PHB and AVA processes in the PHB/AVA CD4 reporter was quantified in Fiji ImageJ (Schindelin *et al*. 2012). Briefly, the entire Z-stack was scanned while tracing over the GFP+ region (where the PHB and AVA processes overlap) with a segmented line and then measuring the overall line length. In cases where the contact and resulting GFP signal was discontinuous, multiple lines were drawn, measured independently, and summed to yield the overall contact site length. For visualization purposes, figures contain a representative subset of Z-stacks reconstructed as maximum intensity projection using Zen Imaging software (blue edition, Zeiss) to display the maximal PHB/AVA contact site.

## SUPPLEMENTAL FIGURE AND TABLE LEGENDS

**Figure S1:**
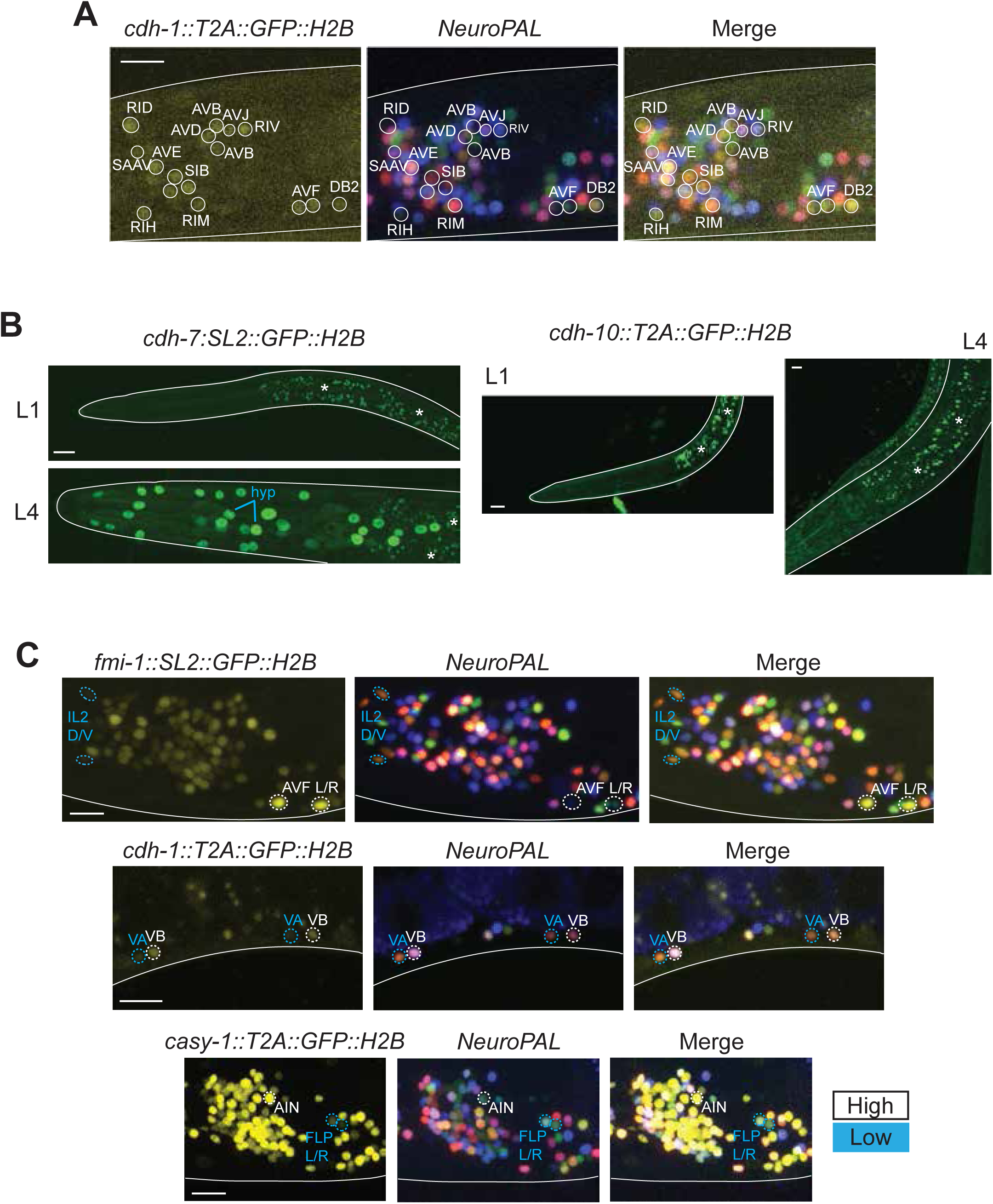
Cellular expression analysis of cadherin reporters. **(A)** Representative images of the *cdh-1::T2A::GFP::H2B* reporter (*ot1092*) showing cell-specific identification of neurons based on overlay with the landmark reporter NeuroPAL (*otIs669*). Images are of L4-stage animals and represent maximum intensity projections of a subset of the Z-stack. **(B)** Representative images of reporters *cdh-7::SL2::GFP::H2B* (*syb4675*) and *cdh-10::T2A::GFP::H2B* (*ot1117*). Gut autofluorescence is marked with an asterisk. **(C)** Differential expression levels for the same cadherin type across neuron types. Representative images of *fmi-1::SL2::GFP::H2B* (*syb4563*), *cdh-1::T2A::GFP::H2B* (*ot1092*), and *casy-1::T2A::GFP::H2B* (*ot1108*) across representative neuron types with low (blue) or high (white) expression.

**Figure S2:**
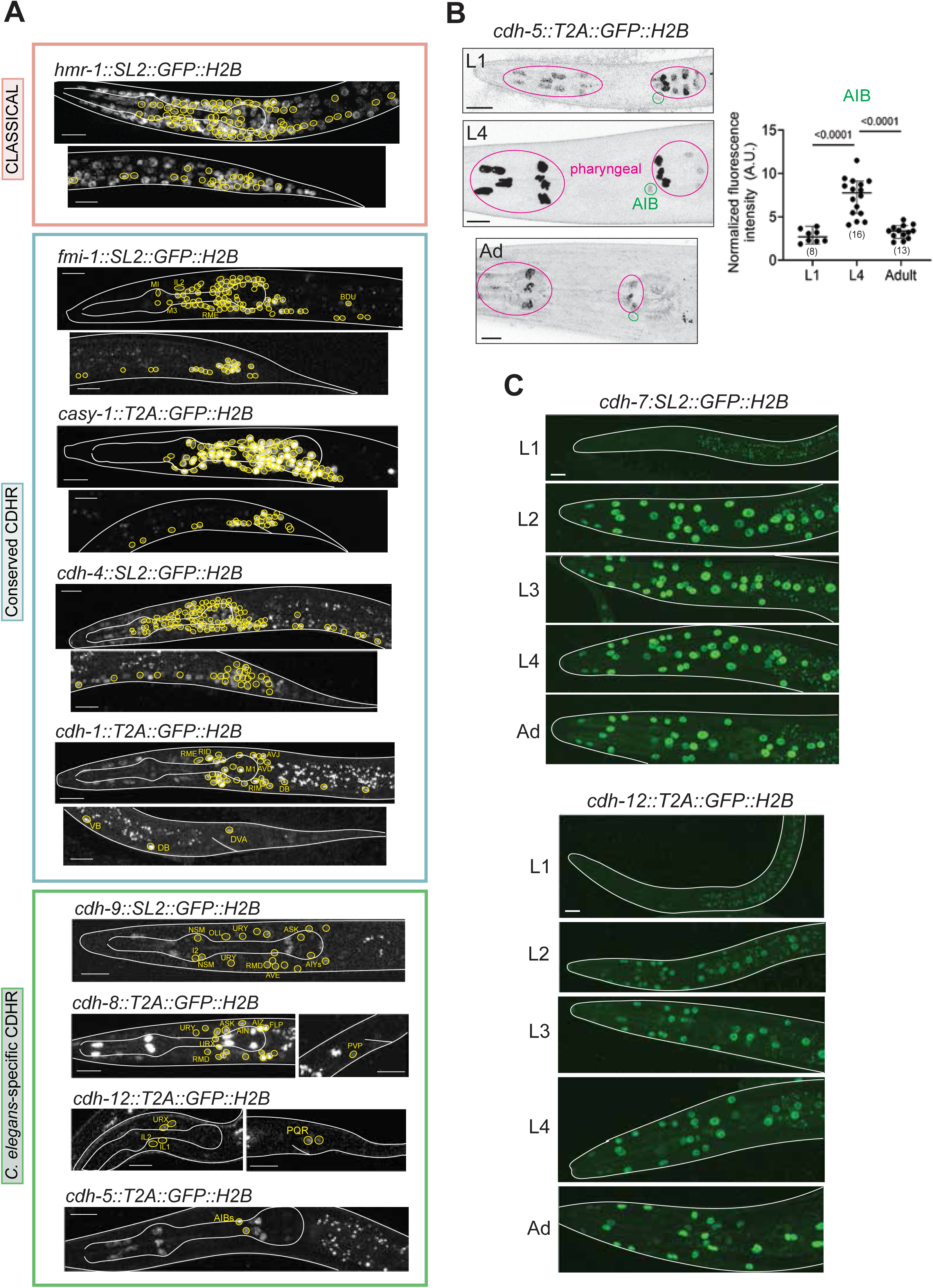
Developmental cadherin expression. **(A)** Representative images of cadherin expression in L1 animals. Reporters of classical cadherin *hmr-1*(*syb4454*) (pink) and non-classical cadherins (green) *casy-1*/*Clstn* (*ot1108*), *cdh-4*/*Fat* (*syb4476*), and *fmi-1*/*Celsr* (*syb4563*) are pan-neuronally or broadly expressed. Reporters of remaining cadherins *cdh-1*(*ot1092*), *cdh-3*(*ot1096*), *cdh-5*(*ot1127*), *cdh-8*(*ot1106*), *cdh-9*(*ot1095*), and *cdh-12*(*ot1119*) are relatively sparsely expressed. All GFP+ neurons are encircled in yellow. Images are of L1-stage animals and represent maximum intensity projections of a subset of the Z-stack. Scale bars = 10μM. **(B)** Developmental changes in *cdh-5(ot1127)* expression. **(C)** *cdh-7*(*syb4675*) and *cdh-12*(*ot1119*) expression onset in the hypodermis begins at L2 and persists until adulthood.

**Figure S3:**
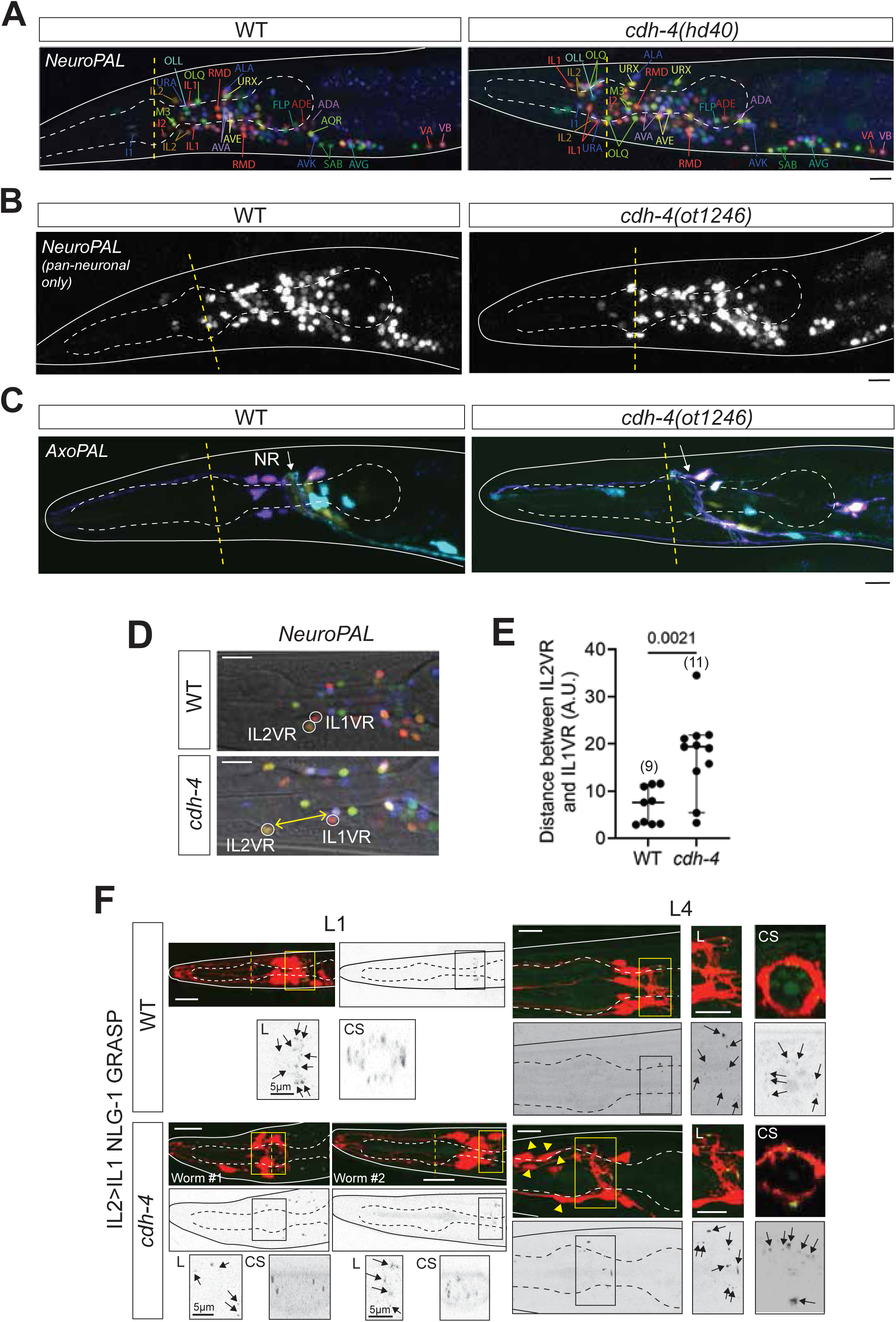
Phenotypic analysis of *cdh-4* mutants. **(A)** Neuronal soma mispositioning defects in *cdh-4*(*hd40*) mutants phenocopy defects in *cdh-4(ot1246)* mutants as assessed with NeuroPAL (*otIs669*). **(B)** Neuron number is unaffected in *cdh-4(ot1246)* mutants, as assessed with a pan-neuronal reporter in the NeuroPAL (*otIs669*) landmark strain. **(C)** Nerve ring anterior shift in *cdh-4(ot1246)* mutants as visualized with AxoPAL (*otEx7895*) reporter. (**D, E**) IL2 and IL1 soma are mispositioned in *cdh-4(ot1246)* mutants. Representative images (D) and quantification (E) of shortest distances (yellow) between IL2VR and IL1VR soma labeled with *NeuroPAL* (*otIs669*) in wild type and *cdh-4* mutant animals at the L4 stage. P-value from unpaired t-test is shown. **(F)** IL2>IL1 synapses labeled with a GRASP reporter (*otIs657*) can still be observed in *cdh-4(hd40)* mutants at L1 and L4 stages. *cdh-4* L1 images show 2 representative worms with variable IL1/IL2 soma mispositioning defects (Worm #1: severe, Worm #2: mild); both populations have GFP puncta despite different number of IL neurons anterior of the midline of anterior pharyngeal bulb (dashed yellow line). L: lateral, CS: cross-section of the nerve ring. P-values from unpaired t-test are shown. 10 animals were analyzed for each panel. Representative images of L4/young adult stage animals are shown and represent maximum intensity projections of the complete Z-stack. Scale bar = 10μM.

**Figure S4:**
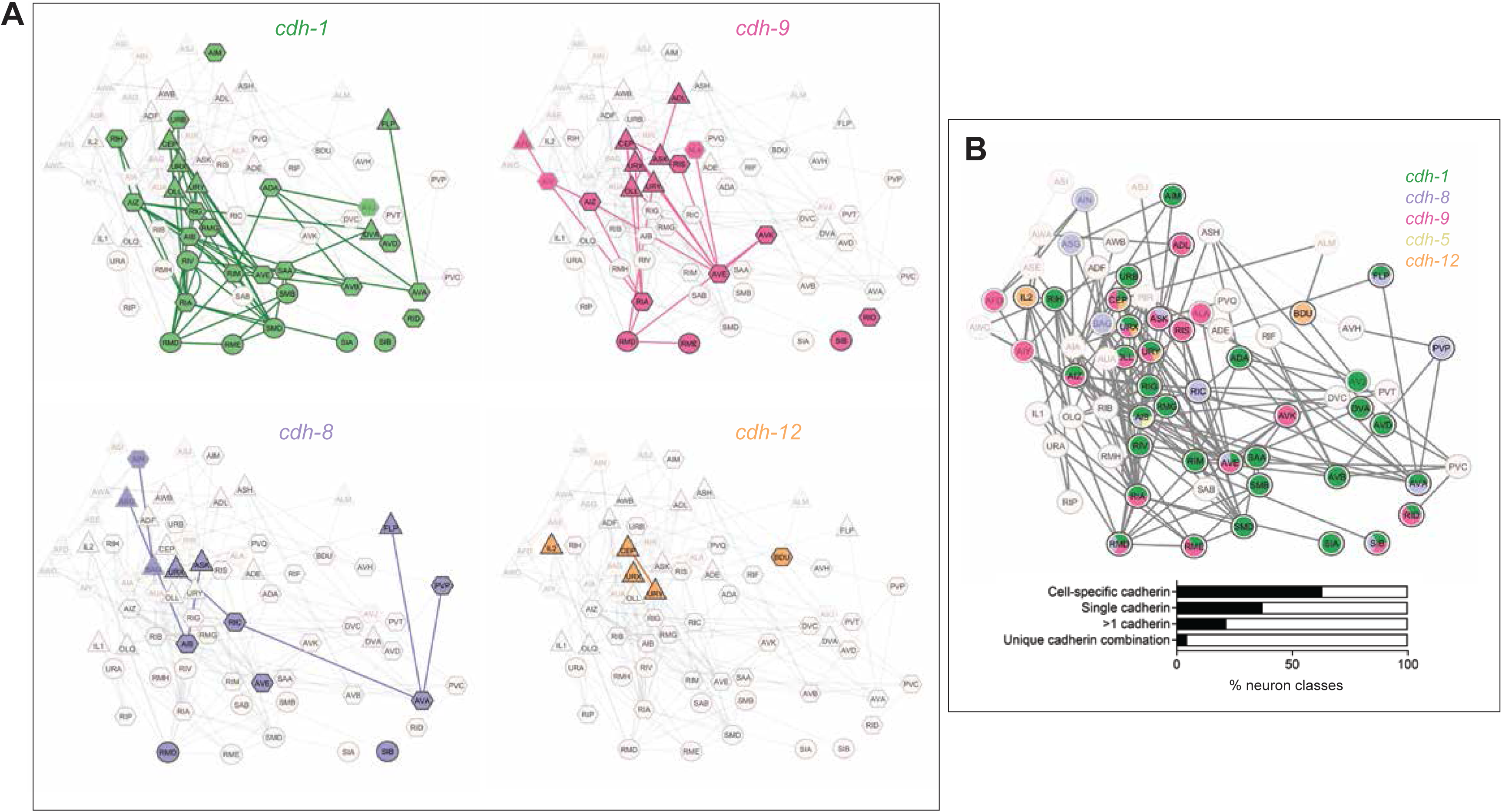
Narrowly expressed cadherins are largely expressed in neurons that have conserved synapses across multiple *C. elegans* samples. **(A)** Expression of *cdh-1*, *cdh-9*, *cdh-8*, *cdh-12*, and *cdh-3* overlaid on the core/conserved *C. elegans* connectome, which includes connectomes for all developmental stages and both sexes (Cook *et al*. 2019; Witvliet *et al*. 2021; Cook *et al*. 2023). Percentages of neurons expressing cell-specific cadherins, single or multiple cadherins, and unique cadherin combinations are shown. **(B)** Expression of all uniquely and narrowly expressed cadherins overlaid on the core connectome.

**Figure S5:**
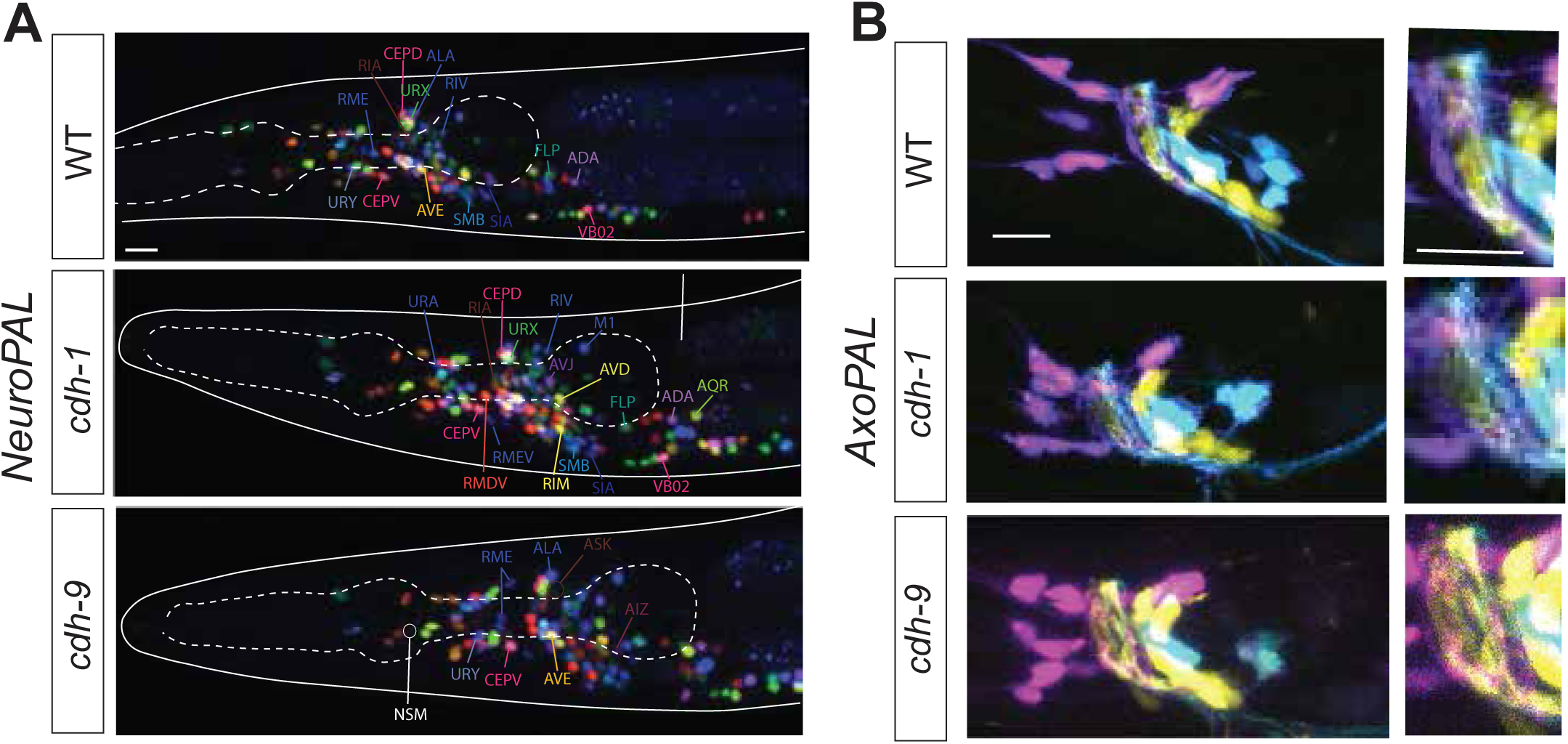
Broad phenotypic analysis of *cdh-1* and *cdh-9* mutants. **(A)** Representative images of *NeuroPAL* reporter in the background of *cdh-1(ot1034)* and *cdh-9(ot1247)* mutants. Only *cdh-1*/*cdh-9* positive neurons are labeled. **(B)** Representative images of AxoPAL reporter in the background of *cdh-1(ot1034)* and *cdh-9(ot1247)* mutants. All images are maximum intensity projections of a subset of the Z-stack. Scale bars = 10μM. 10 animals per genotype were analyzed for the NeuroPAL and AxoPAL analyses.

**Figure S6:**
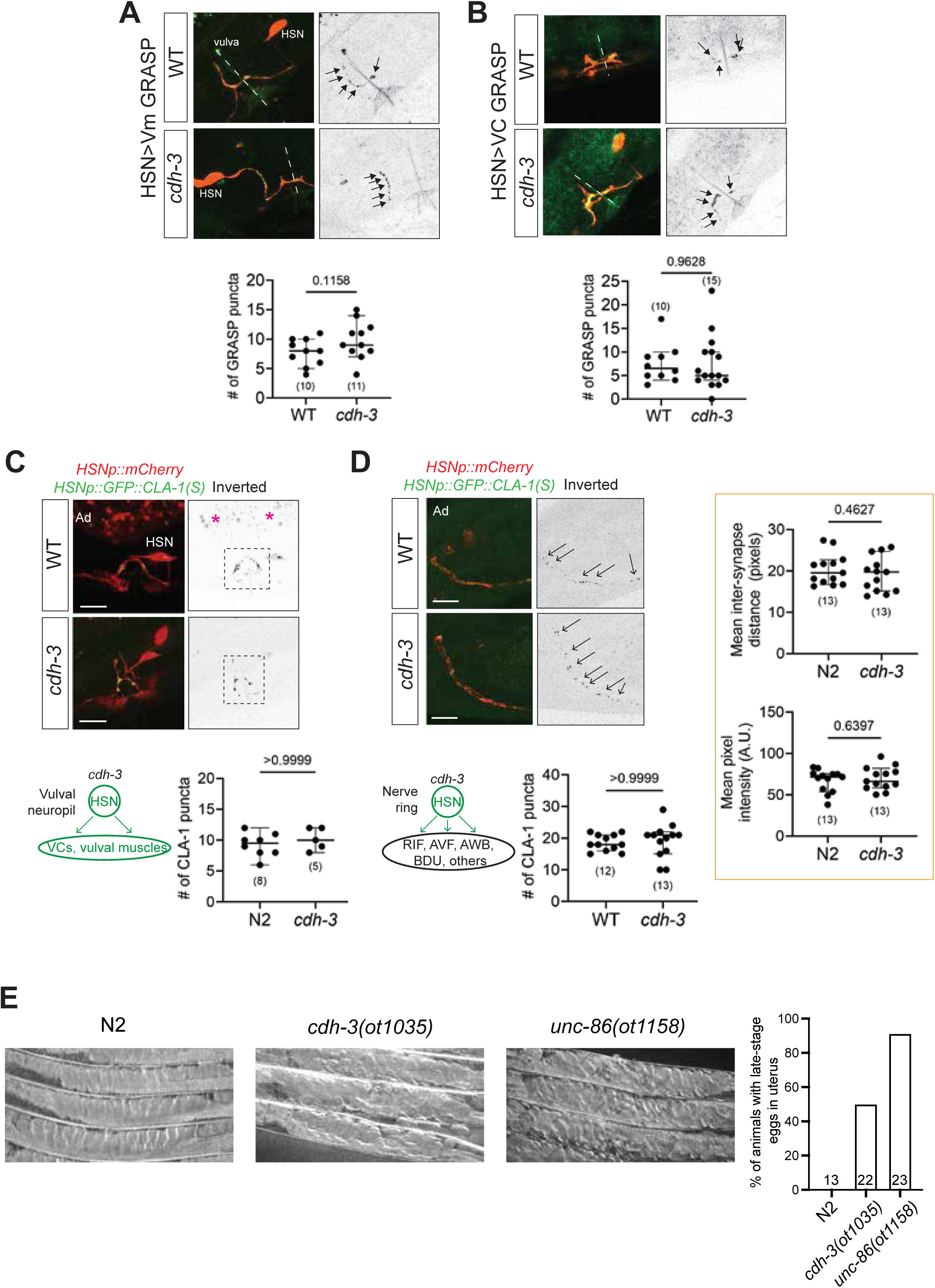
Phenotypic analysis of *cdh-3/Fat* mutants. **(A)** HSN>Vm GRASP signal is unaffected in *cdh-3(ot1035)* mutants. **(B)** HSN>VC GRASP signal is unaffected in *cdh-3(ot1035)* mutants. **(C, D)** HSN-specific presynaptic CLA-1::GFP puncta in the vulval region (C) and nerve ring (D) are unaffected in *cdh-3(ot1035)* mutants. Quantitative features other than puncta number – scored using WormPsyQi - are also unaffected in the nerve ring. Gut autofluorescence is marked with an asterisk. **(E)** Egg-laying defects in *cdh-3(ot1035)* mutants were compared to wild-type N2 and *unc-86(ot1158)* null mutants (positive control). Late-stage eggs are embryos that have passed the comma stage (430 min of development). Eggs are normally laid at ∼150 min of embryonic development. All images are maximum intensity projections of a subset of the Z-stack. Scale bars = 10μM. In all graphs, a dot represents one worm and error bars denote median with 95% confidence interval. P-values from unpaired t-test are shown.

**Figure S7:**
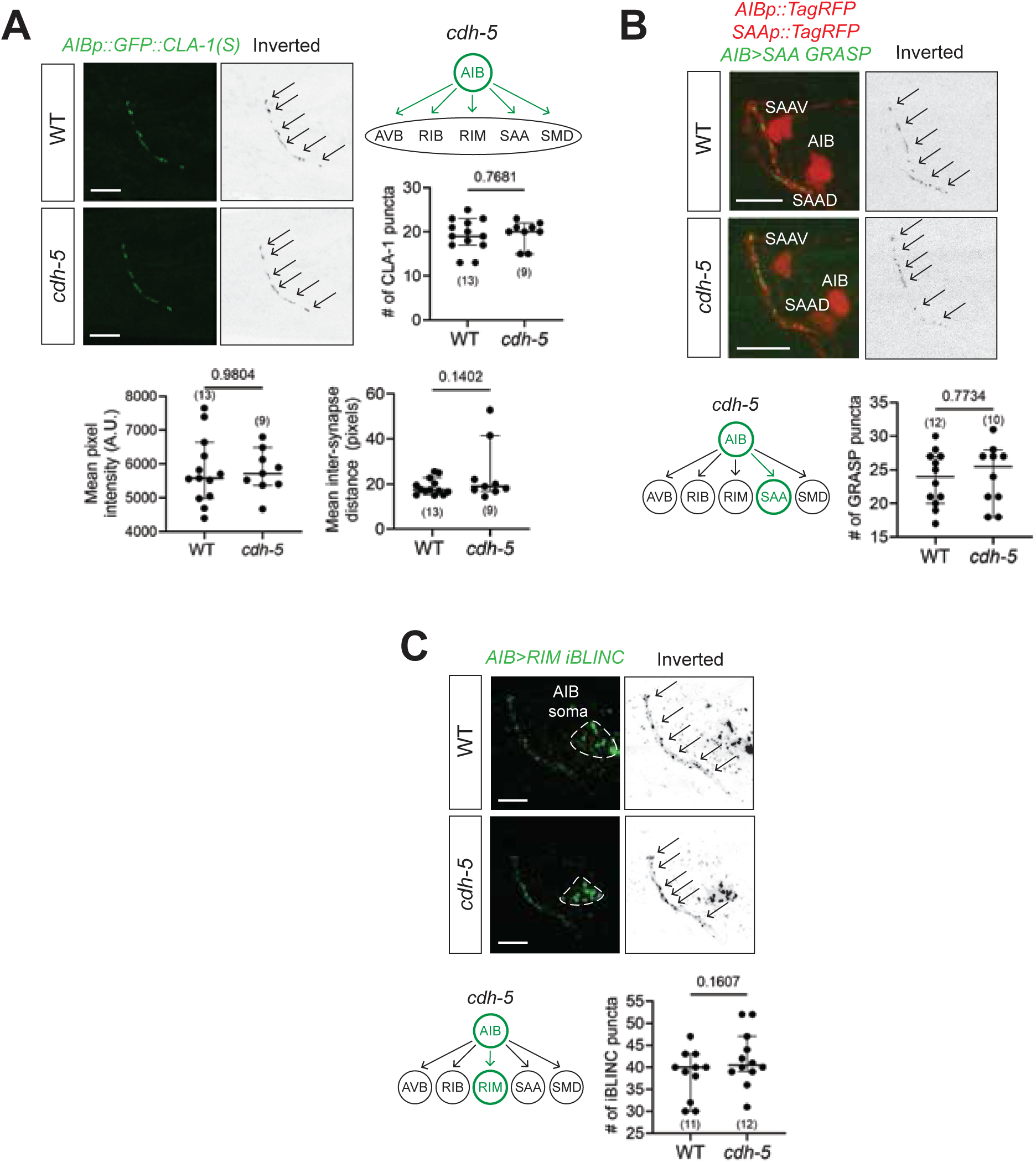
Chemical synapses are unaffected in *cdh-5* mutants. **(A)** AIB presynaptic specializations visualized with *AIBp::GFP::CLA-1* (*otIs886*) are unaffected in *cdh-5(ot1093)* null mutants. *cdh-5* is exclusively expressed in AIB neurons. No phenotype was observed in the number, mean pixel intensity, and mean inter-synaptic distance of CLA-1 puncta. **(B)** AIB>SAA synapses, assessed with an NLG-1 GRASP reporter (*otEx7809*), are unaffected in *cdh-5(ot1093)* mutants. *cdh-5* is expressed only in AIB neurons. **(C)** AIB>RIM synapses assessed with an iBLINC reporter (*dzIs89*) are unaffected in *cdh-5(ot1093)* mutants. *cdh-5* is expressed only in AIB neurons. All images are maximum intensity projections of a subset of the Z-stack. Scale bars = 10μM. In all graphs, a dot represents one worm and error bars denote median with 95% confidence interval. P-values from unpaired t-test are shown.

**Table S1:**
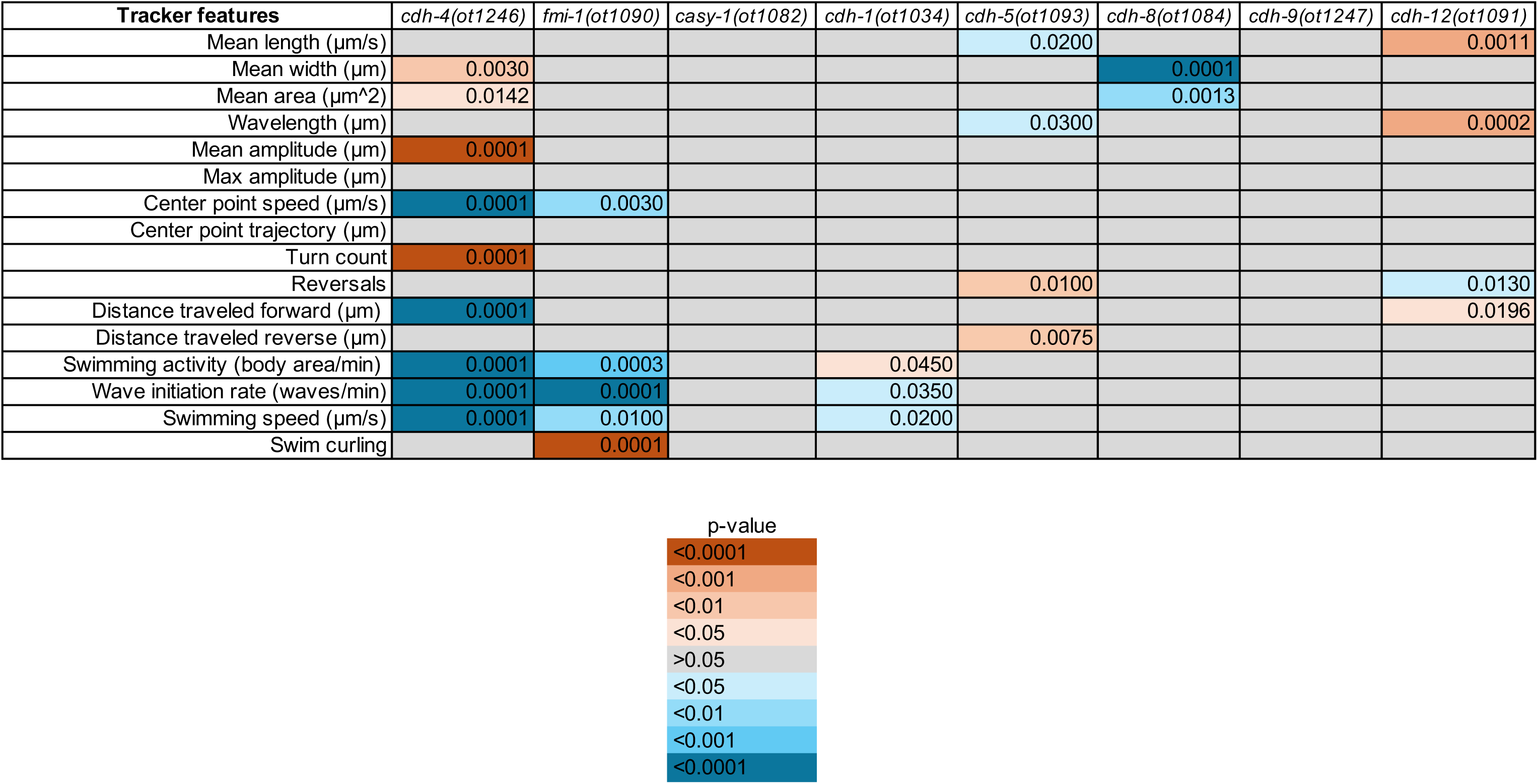
Wormtracker data.

**Table S2:**
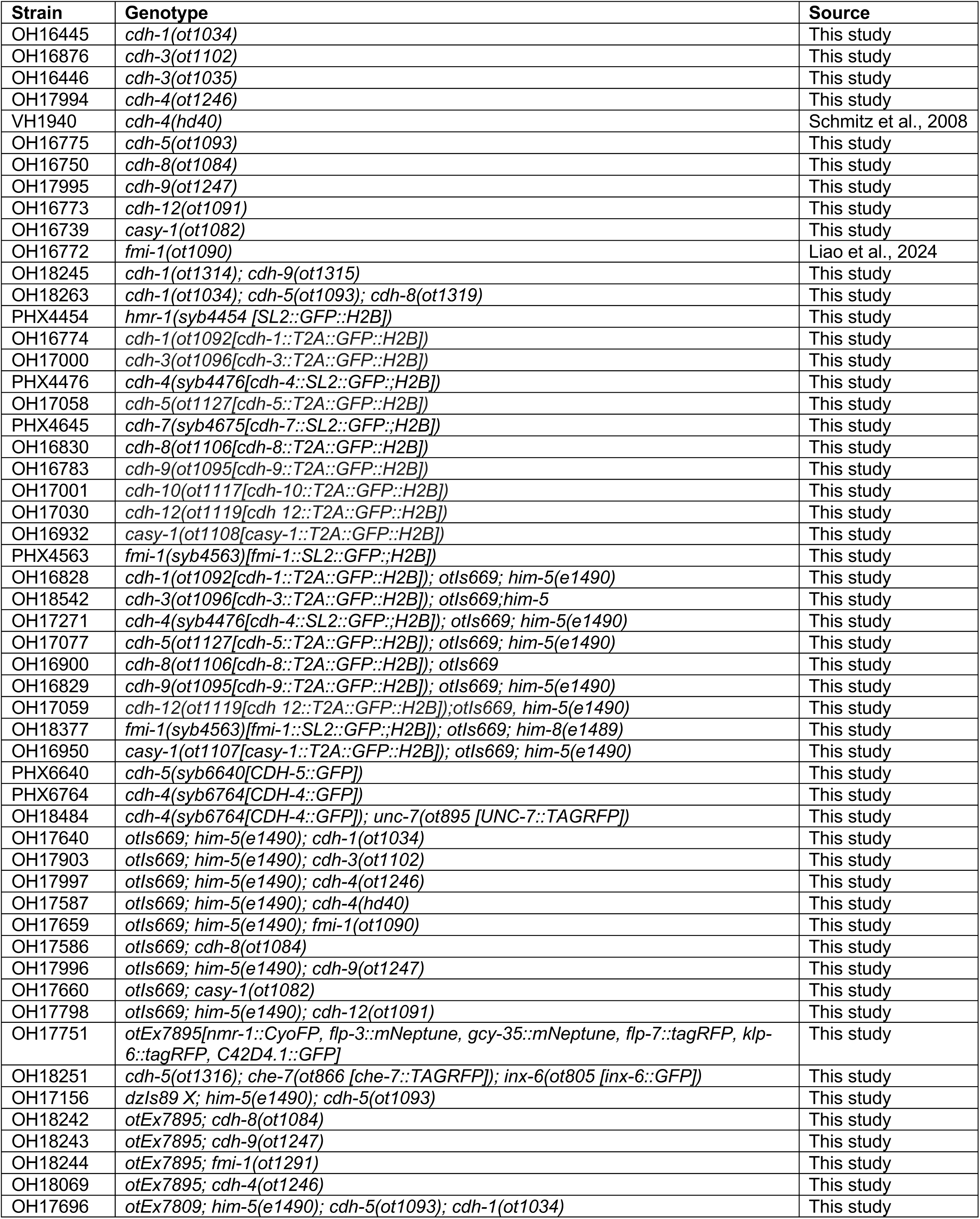
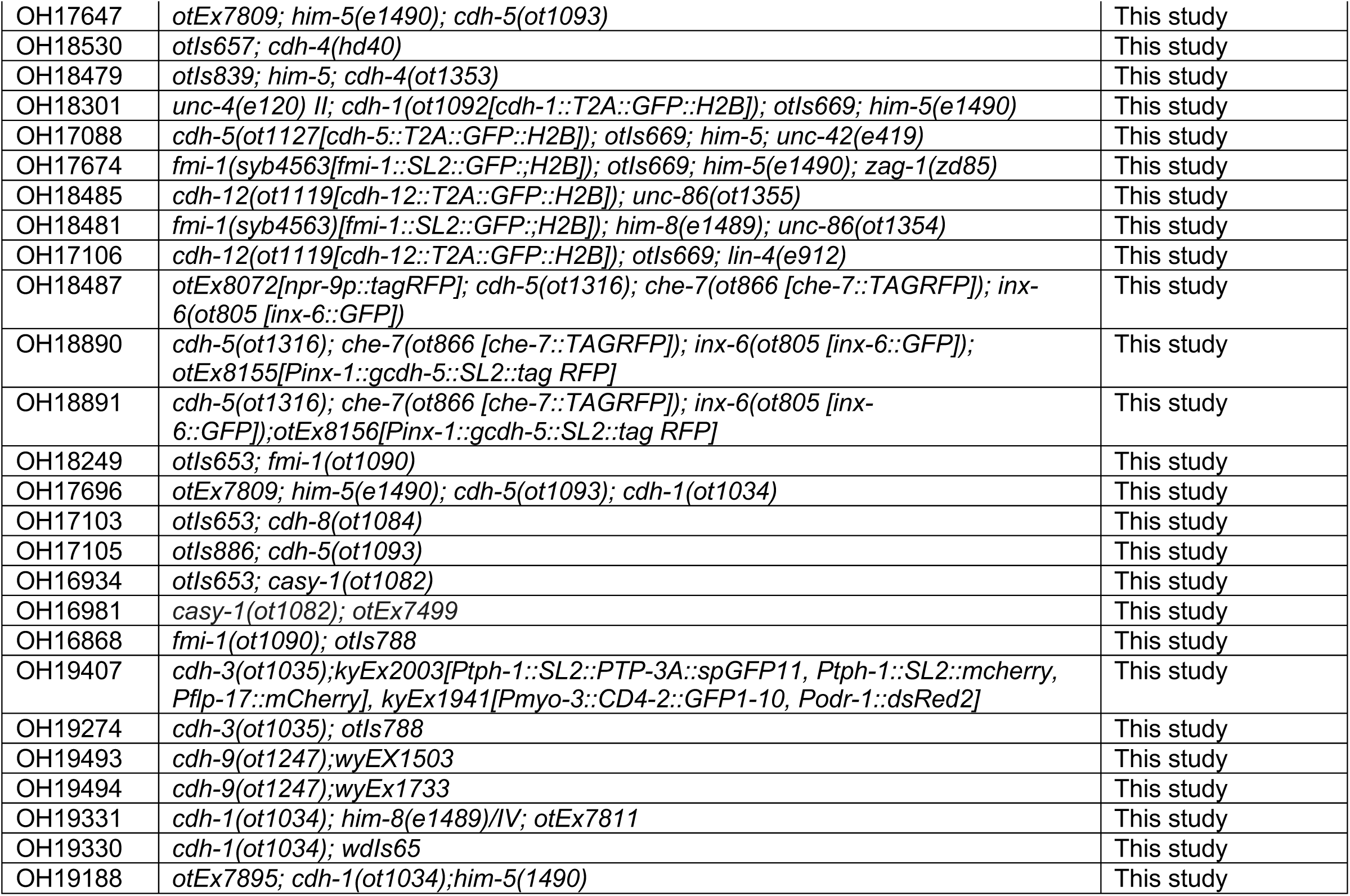
Strain list.

## Notes

### Competing Interest Statement

The authors have declared no competing interest.

## REFERENCES

Abedin, M., and N. King, 2008 The premetazoan ancestry of cadherins. Science 319: 946–948.

Albertin, C. B., O. Simakov, T. Mitros, Z. Y. Wang, J. R. Pungor et al., 2015 The octopus genome and the evolution of cephalopod neural and morphological novelties. Nature 524: 220–224.

Ambros, V., and H. R. Horvitz, 1984 Heterochronic mutants of the nematode Caenorhabditis elegans. Science 226: 409–416.

Aviles, E. C., and L. V. Goodrich, 2017 Configuring a robust nervous system with Fat cadherins. Semin Cell Dev Biol 69: 91–101.

Barnes, K. M., L. Fan, M. W. Moyle, C. A. Brittin, Y. Xu et al., 2020 Cadherin preserves cohesion across involuting tissues during C. elegans neurulation. Elife 9.

Basu, R., X. Duan, M. R. Taylor, E. A. Martin, S. Muralidhar et al., 2017 Heterophilic Type II Cadherins Are Required for High-Magnitude Synaptic Potentiation in the Hippocampus. Neuron 96: 160–176 e168.

Bayer, E., and O. Hobert, 2018 A novel null allele of C. elegans gene ceh-14. MicroPubl Biol 2018.

Berghoff, E. G., L. Glenwinkel, A. Bhattacharya, H. Sun, E. Varol et al., 2021 The Prop1-like homeobox gene unc-42 specifies the identity of synaptically connected neurons. Elife 10.

Bhattacharya, A., U. Aghayeva, E. G. Berghoff and O. Hobert, 2019 Plasticity of the Electrical Connectome of C. elegans. Cell 176: 1174–1189 e1116.

Bischoff, M., and R. Schnabel, 2006 Global cell sorting is mediated by local cell-cell interactions in the C. elegans embryo. Dev Biol 294: 432–444.

Brasch, J., O. J. Harrison, B. Honig and L. Shapiro, 2012 Thinking outside the cell: how cadherins drive adhesion. Trends Cell Biol 22: 299–310.

Brasch, J., P. S. Katsamba, O. J. Harrison, G. Ahlsen, R. B. Troyanovsky et al., 2018 Homophilic and Heterophilic Interactions of Type II Cadherins Identify Specificity Groups Underlying Cell-Adhesive Behavior. Cell Rep 23: 1840–1852.

Brenner, S., 1974 The genetics of Caenorhabditis elegans. Genetics 77: 71–94.

Brittin, C. A., S. J. Cook, D. H. Hall, S. W. Emmons and N. Cohen, 2021 A multi-scale brain map derived from whole-brain volumetric reconstructions. Nature 591: 105–110.

Broadbent, I. D., and J. Pettitt, 2002 The C. elegans hmr-1 gene can encode a neuronal classic cadherin involved in the regulation of axon fasciculation. Curr Biol 12: 59–63.

Cappello, S., M. J. Gray, C. Badouel, S. Lange, M. Einsiedler et al., 2013 Mutations in genes encoding the cadherin receptor-ligand pair DCHS1 and FAT4 disrupt cerebral cortical development. Nat Genet 45: 1300–1308.

Cardenas-Garcia, S. P., S. Ijaz and A. E. Pereda, 2023 The components of an electrical synapse as revealed by expansion microscopy of a single synaptic contact. bioRxiv.

Cassada, R. C., and R. L. Russell, 1975 The dauerlarva, a post-embryonic developmental variant of the nematode Caenorhabditis elegans. Dev Biol 46: 326–342.

Chai, G., L. Zhou, M. Manto, F. Helmbacher, F. Clotman et al., 2014 Celsr3 is required in motor neurons to steer their axons in the hindlimb. Nat Neurosci 17: 1171–1179.

Chen, P. L., and T. R. Clandinin, 2008 The cadherin Flamingo mediates level-dependent interactions that guide photoreceptor target choice in Drosophila. Neuron 58: 26–33.

Clark, S. G., and C. Chiu, 2003 C. elegans ZAG-1, a Zn-finger-homeodomain protein, regulates axonal development and neuronal differentiation. Development 130: 3781–3794.

Cook, S. J., C. M. Crouse, E. Yemini, D. H. Hall, S. W. Emmons et al., 2020 The connectome of the Caenorhabditis elegans pharynx. J Comp Neurol 528: 2767–2784.

Cook, S. J., T. A. Jarrell, C. A. Brittin, Y. Wang, A. E. Bloniarz et al., 2019 Whole-animal connectomes of both Caenorhabditis elegans sexes. Nature 571: 63–71.

Cook, S. J., C. A. Kalinski and O. Hobert, 2023 Neuronal contact predicts connectivity in the C. elegans brain. Curr Biol 33: 2315–2320 e2312.

Costa, M., W. Raich, C. Agbunag, B. Leung, J. Hardin et al., 1998 A putative catenin-cadherin system mediates morphogenesis of the Caenorhabditis elegans embryo. J Cell Biol 141: 297–308.

Cros, C., and O. Hobert, 2022 Caenorhabditis elegans sine oculis/SIX-type homeobox genes act as homeotic switches to define neuronal subtype identities. Proc Natl Acad Sci U S A 119: e2206817119.

de Wit, J., and A. Ghosh, 2016 Specification of synaptic connectivity by cell surface interactions. Nat Rev Neurosci 17: 22–35.

Dempsey, W. P., S. E. Fraser and P. Pantazis, 2012 PhOTO zebrafish: a transgenic resource for in vivo lineage tracing during development and regeneration. PLoS One 7: e32888.

Desbois, M., S. J. Cook, S. W. Emmons and H. E. Bulow, 2015 Directional Trans-Synaptic Labeling of Specific Neuronal Connections in Live Animals. Genetics 200: 697–705.

Duan, X., A. Krishnaswamy, I. De la Huerta and J. R. Sanes, 2014 Type II cadherins guide assembly of a direction-selective retinal circuit. Cell 158: 793–807.

Duan, X., A. Krishnaswamy, M. A. Laboulaye, J. Liu, Y. R. Peng et al., 2018 Cadherin Combinations Recruit Dendrites of Distinct Retinal Neurons to a Shared Interneuronal Scaffold. Neuron 99: 1145–1154 e1146.

Fan, L., I. Kovacevic, M. G. Heiman and Z. Bao, 2019 A multicellular rosette-mediated collective dendrite extension. Elife 8.

Feinberg, E. H., M. K. Vanhoven, A. Bendesky, G. Wang, R. D. Fetter et al., 2008 GFP Reconstitution Across Synaptic Partners (GRASP) defines cell contacts and synapses in living nervous systems. Neuron 57: 353–363.

Friedman, L. G., D. L. Benson and G. W. Huntley, 2015 Cadherin-based transsynaptic networks in establishing and modifying neural connectivity. Curr Top Dev Biol 112: 415–465.

Fulford, A. D., and H. McNeill, 2020 Fat/Dachsous family cadherins in cell and tissue organisation. Curr Opin Cell Biol 62: 96–103.

Furness, J. B., and M. J. Stebbing, 2018 The first brain: Species comparisons and evolutionary implications for the enteric and central nervous systems. Neurogastroenterol Motil 30.

Ghanta, K. S., T. Ishidate and C. C. Mello, 2021 Microinjection for precision genome editing in Caenorhabditis elegans. STAR Protoc 2: 100748.

Hill, E., I. D. Broadbent, C. Chothia and J. Pettitt, 2001 Cadherin superfamily proteins in Caenorhabditis elegans and Drosophila melanogaster. J Mol Biol 305: 1011–1024.

Hirano, S., and M. Takeichi, 2012 Cadherins in brain morphogenesis and wiring. Physiol Rev 92: 597–634.

Hobert, O., 2016 Terminal Selectors of Neuronal Identity. Curr Top Dev Biol 116: 455–475.

Hoerndli, F. J., M. Walser, E. Frohli Hoier, D. de Quervain, A. Papassotiropoulos et al., 2009 A conserved function of C. elegans CASY-1 calsyntenin in associative learning. PLoS One 4: e4880.

Honig, B., and L. Shapiro, 2020 Adhesion Protein Structure, Molecular Affinities, and Principles of Cell-Cell Recognition. Cell 181: 520–535.

Hsu, H. W., C. P. Liao, Y. C. Chiang, R. T. Syu and C. L. Pan, 2020 Caenorhabditis elegans Flamingo FMI-1 controls dendrite self-avoidance through F-actin assembly. Development 147.

Hu, Y., I. Flockhart, A. Vinayagam, C. Bergwitz, B. Berger et al., 2011 An integrative approach to ortholog prediction for disease-focused and other functional studies. BMC Bioinformatics 12: 357.

Hulpiau, P., and F. van Roy, 2011 New insights into the evolution of metazoan cadherins. Mol Biol Evol 28: 647–657.

Hutter, H., 2003 Extracellular cues and pioneers act together to guide axons in the ventral cord of C. elegans. Development 130: 5307–5318.

Hynes, R. O., and Q. Zhao, 2000 The evolution of cell adhesion. J Cell Biol 150: F89–96.

Ikeda, D. D., Y. Duan, M. Matsuki, H. Kunitomo, H. Hutter et al., 2008 CASY-1, an ortholog of calsyntenins/alcadeins, is essential for learning in Caenorhabditis elegans. Proc Natl Acad Sci U S A 105: 5260–5265.

Jain, S., Y. Lin, Y. Z. Kurmangaliyev, J. Valdes-Aleman, S. A. LoCascio et al., 2022 A global timing mechanism regulates cell-type-specific wiring programmes. Nature 603: 112–118.

Jin, E. J., S. Park, X. Lyu and Y. Jin, 2020 Gap junctions: historical discoveries and new findings in the Caenorhabditiselegans nervous system. Biol Open 9.

Jumper, J., R. Evans, A. Pritzel, T. Green, M. Figurnov et al., 2021 Highly accurate protein structure prediction with AlphaFold. Nature 596: 583–589.

Kerk, S. Y., P. Kratsios, M. Hart, R. Mourao and O. Hobert, 2017 Diversification of C. elegans Motor Neuron Identity via Selective Effector Gene Repression. Neuron 93: 80–98.

Kim, B., and S. W. Emmons, 2017 Multiple conserved cell adhesion protein interactions mediate neural wiring of a sensory circuit in C. elegans. Elife 6.

Knufer, A., G. Diana, G. S. Walsh, J. D. Clarke and S. Guthrie, 2020 Cadherins regulate nuclear topography and function of developing ocular motor circuitry. Elife 9.

Kurusu, M., T. Katsuki, K. Zinn and E. Suzuki, 2012 Developmental changes in expression, subcellular distribution, and function of Drosophila N-cadherin, guided by a cell-intrinsic program during neuronal differentiation. Dev Biol 366: 204–217.

Kuwako, K. I., Y. Nishimoto, S. Kawase, H. J. Okano and H. Okano, 2014 Cadherin-7 regulates mossy fiber connectivity in the cerebellum. Cell Rep 9: 311–323.

Liao, C. P., M. Majeed and O. Hobert, 2024 Experience-dependent, sexually dimorphic synaptic connectivity defined by sex-specific cadherin expression. bioRxiv.

Loveless, T., and J. Hardin, 2012 Cadherin complexity: recent insights into cadherin superfamily function in C. elegans. Curr Opin Cell Biol 24: 695–701.

Lv, X., S. Li, J. Li, X. Y. Yu, X. Ge et al., 2022 Patterned cPCDH expression regulates the fine organization of the neocortex. Nature 612: 503–511.

Majeed, M., H. Han, K. Zhang, W. X. Cao, C. P. Liao et al., 2024 Toolkits for detailed and high-throughput interrogation of synapses in C. elegans. Elife 12.

Mangione, F., and E. Martin-Blanco, 2018 The Dachsous/Fat/Four-Jointed Pathway Directs the Uniform Axial Orientation of Epithelial Cells in the Drosophila Abdomen. Cell Rep 25: 2836–2850 e2834.

Miller, D. M., M. M. Shen, C. E. Shamu, T. R. Burglin, G. Ruvkun et al., 1992 C. elegans unc-4 gene encodes a homeodomain protein that determines the pattern of synaptic input to specific motor neurons. Nature 355: 841–845.

Mountoufaris, G., D. Canzio, C. L. Nwakeze, W. V. Chen and T. Maniatis, 2018 Writing, Reading, and Translating the Clustered Protocadherin Cell Surface Recognition Code for Neural Circuit Assembly. Annu Rev Cell Dev Biol 34: 471–493.

Mountoufaris, G., W. V. Chen, Y. Hirabayashi, S. O’Keeffe, M. Chevee et al., 2017 Multicluster Pcdh diversity is required for mouse olfactory neural circuit assembly. Science 356: 411–414.

Moyle, M. W., K. M. Barnes, M. Kuchroo, A. Gonopolskiy, L. H. Duncan et al., 2021 Structural and developmental principles of neuropil assembly in C. elegans. Nature 591: 99–104.

Najarro, E. H., L. Wong, M. Zhen, E. P. Carpio, A. Goncharov et al., 2012 Caenorhabditis elegans Flamingo Cadherin fmi-1 Regulates GABAergic Neuronal Development. J Neurosci 32: 4196–4211.

Nichols, S. A., B. W. Roberts, D. J. Richter, S. R. Fairclough and N. King, 2012 Origin of metazoan cadherin diversity and the antiquity of the classical cadherin/beta-catenin complex. Proc Natl Acad Sci U S A 109: 13046–13051.

Nollet, F., P. Kools and F. van Roy, 2000 Phylogenetic analysis of the cadherin superfamily allows identification of six major subfamilies besides several solitary members. J Mol Biol 299: 551–572.

Ohno, H., S. Kato, Y. Naito, H. Kunitomo, M. Tomioka et al., 2014 Role of synaptic phosphatidylinositol 3-kinase in a behavioral learning response in C. elegans. Science 345: 313–317.

Packer, J. S., Q. Zhu, C. Huynh, P. Sivaramakrishnan, E. Preston et al., 2019 A lineage-resolved molecular atlas of C. elegans embryogenesis at single-cell resolution. Science 365.

Pereira, L., F. Aeschimann, C. Wang, H. Lawson, E. Serrano-Saiz et al., 2019 Timing mechanism of sexually dimorphic nervous system differentiation. Elife 8.

Pettitt, J., W. B. Wood and R. H. Plasterk, 1996 cdh-3, a gene encoding a member of the cadherin superfamily, functions in epithelial cell morphogenesis in Caenorhabditis elegans. Development 122: 4149–4157.

Pham, K., N. Masoudi, E. Leyva-Diaz and O. Hobert, 2021 A nervous system-specific subnuclear organelle in Caenorhabditis elegans. Genetics 217: 1–17.

Polanco, J., F. Reyes-Vigil, S. D. Weisberg, I. Dhimitruka and J. L. Bruses, 2021 Differential Spatiotemporal Expression of Type I and Type II Cadherins Associated With the Segmentation of the Central Nervous System and Formation of Brain Nuclei in the Developing Mouse. Front Mol Neurosci 14: 633719.

Quevillon, E., V. Silventoinen, S. Pillai, N. Harte, N. Mulder et al., 2005 InterProScan: protein domains identifier. Nucleic Acids Res 33: W116–120.

Reilly, M. B., C. Cros, E. Varol, E. Yemini and O. Hobert, 2020 Unique homeobox codes delineate all the neuron classes of C. elegans. Nature 584: 595–601.

Restif, C., C. Ibanez-Ventoso, M. M. Vora, S. Guo, D. Metaxas et al., 2014 CeleST: computer vision software for quantitative analysis of C. elegans swim behavior reveals novel features of locomotion. PLoS Comput Biol 10: e1003702.

Ripoll-Sanchez, L., J. Watteyne, H. Sun, R. Fernandez, S. R. Taylor et al., 2023 The neuropeptidergic connectome of C. elegans. Neuron 111: 3570–3589 e3575.

Sanes, J. R., and S. L. Zipursky, 2020 Synaptic Specificity, Recognition Molecules, and Assembly of Neural Circuits. Cell 181: 536–556.

Schafer, W. R., 2005 Egg-laying. WormBook: 1–7.

Schindelin, J., I. Arganda-Carreras, E. Frise, V. Kaynig, M. Longair et al., 2012 Fiji: an open-source platform for biological-image analysis. Nat Methods 9: 676–682.

Schmitz, C., I. Wacker and H. Hutter, 2008 The Fat-like cadherin CDH-4 controls axon fasciculation, cell migration and hypodermis and pharynx development in Caenorhabditis elegans. Dev Biol 316: 249–259.

Schnabel, R., M. Bischoff, A. Hintze, A. K. Schulz, A. Hejnol et al., 2006 Global cell sorting in the C. elegans embryo defines a new mechanism for pattern formation. Dev Biol 294: 418–431.

Schwabe, T., J. A. Borycz, I. A. Meinertzhagen and T. R. Clandinin, 2014 Differential adhesion determines the organization of synaptic fascicles in the Drosophila visual system. Curr Biol 24: 1304–1313.

Schwabe, T., H. Neuert and T. R. Clandinin, 2013 A network of cadherin-mediated interactions polarizes growth cones to determine targeting specificity. Cell 154: 351–364.

Serrano-Saiz, E., Richard J. Poole, T. Felton, F. Zhang, Estanisla D. De La Cruz et al., 2013 Modular Control of Glutamatergic Neuronal Identity in C. elegans by Distinct Homeodomain Proteins. Cell 155: 659–673.

Shaham, S., and C. I. Bargmann, 2002 Control of neuronal subtype identity by the C. elegans ARID protein CFI-1. Genes Dev 16: 972–983.

Shapiro, L., A. M. Fannon, P. D. Kwong, A. Thompson, M. S. Lehmann et al., 1995a Structural basis of cell-cell adhesion by cadherins. Nature 374: 327–337.

Shapiro, L., P. D. Kwong, A. M. Fannon, D. R. Colman and W. A. Hendrickson, 1995b Considerations on the folding topology and evolutionary origin of cadherin domains. Proc Natl Acad Sci U S A 92: 6793–6797.

Shen, K., and C. I. Bargmann, 2003 The immunoglobulin superfamily protein SYG-1 determines the location of specific synapses in C. elegans. Cell 112: 619–630.

Steimel, A., L. Wong, E. H. Najarro, B. D. Ackley, G. Garriga et al., 2010 The Flamingo ortholog FMI-1 controls pioneer-dependent navigation of follower axons in C. elegans. Development 137: 3663–3673.

Sun, H., and O. Hobert, 2021 Temporal transitions in the post-mitotic nervous system of Caenorhabditis elegans. Nature 600: 93–99.

Sun, Y. C., X. Chen, S. Fischer, S. Lu, H. Zhan et al., 2021 Integrating barcoded neuroanatomy with spatial transcriptional profiling enables identification of gene correlates of projections. Nat Neurosci 24: 873–885.

Sundararajan, L., M. L. Norris, S. Schoneich, B. D. Ackley and E. A. Lundquist, 2014 The fat-like cadherin CDH-4 acts cell-non-autonomously in anterior-posterior neuroblast migration. Dev Biol 392: 141–152.

Takeichi, M., 1988 The cadherins: cell-cell adhesion molecules controlling animal morphogenesis. Development 102: 639–655.

Taylor, S. R., G. Santpere, A. Weinreb, A. Barrett, M. B. Reilly et al., 2021 Molecular topography of an entire nervous system. Cell 184: 4329–4347 e4323.

Thakar, S., L. Wang, T. Yu, M. Ye, K. Onishi et al., 2017 Evidence for opposing roles of Celsr3 and Vangl2 in glutamatergic synapse formation. Proc Natl Acad Sci U S A 114: E610–E618.

Thapliyal, S., S. Ravindranath and K. Babu, 2018a Regulation of Glutamate Signaling in the Sensorimotor Circuit by CASY-1A/Calsyntenin in Caenorhabditis elegans. Genetics 208: 1553–1564.

Thapliyal, S., A. Vasudevan, Y. Dong, J. Bai, S. P. Koushika et al., 2018b The C-terminal of CASY-1/Calsyntenin regulates GABAergic synaptic transmission at the Caenorhabditis elegans neuromuscular junction. PLoS Genet 14: e1007263.

Tsai, T. Y., M. Sikora, P. Xia, T. Colak-Champollion, H. Knaut et al., 2020 An adhesion code ensures robust pattern formation during tissue morphogenesis. Science 370: 113–116.

Vidal, B., B. Gulez, W. X. Cao, E. Leyva-Diaz, M. B. Reilly et al., 2022 The enteric nervous system of the C. elegans pharynx is specified by the Sine oculis-like homeobox gene ceh-34. Elife 11.

Wacker, I., V. Schwarz, E. M. Hedgecock and H. Hutter, 2003 zag-1, a Zn-finger homeodomain transcription factor controlling neuronal differentiation and axon outgrowth in C. elegans. Development 130: 3795–3805.

Wang, J., H. T. Schwartz and M. M. Barr, 2010 Functional specialization of sensory cilia by an RFX transcription factor isoform. Genetics 186: 1295–1307.

Ward, S., N. Thomson, J. G. White and S. Brenner, 1975 Electron microscopical reconstruction of the anterior sensory anatomy of the nematode Caenorhabditis elegans.?2UU. J Comp Neurol 160: 313–337.

White, J. G., E. Southgate, J. N. Thomson and S. Brenner, 1986 The structure of the nervous system of the nematode Caenorhabditis elegans. Philos Trans R Soc Lond B Biol Sci 314: 1–340.

Williams, M. E., S. A. Wilke, A. Daggett, E. Davis, S. Otto et al., 2011 Cadherin-9 regulates synapse-specific differentiation in the developing hippocampus. Neuron 71: 640–655.

Witvliet, D., B. Mulcahy, J. K. Mitchell, Y. Meirovitch, D. R. Berger et al., 2021 Connectomes across development reveal principles of brain maturation. Nature 596: 257–261.

Wolterhoff, N., and P. R. Hiesinger, 2024 Synaptic promiscuity in brain development. Curr Biol 34: R102–R116.

Yamagata, M., X. Duan and J. R. Sanes, 2018 Cadherins Interact With Synaptic Organizers to Promote Synaptic Differentiation. Front Mol Neurosci 11: 142.

Yemini, E., A. Lin, A. Nejatbakhsh, E. Varol, R. Sun et al., 2021 NeuroPAL: A Multicolor Atlas for Whole-Brain Neuronal Identification in C. elegans. Cell 184: 272–288 e211.

Zipursky, S. L., and J. R. Sanes, 2010 Chemoaffinity revisited: dscams, protocadherins, and neural circuit assembly. Cell 143: 343–353.

Zou, Y., 2020 Breaking symmetry - cell polarity signaling pathways in growth cone guidance and synapse formation. Curr Opin Neurobiol 63: 77–86.

